# The Hard Limits of Decoding Mental States: The Decodability of fMRI

**DOI:** 10.1101/2020.12.18.423495

**Authors:** R. Jabakhanji, A.D. Vigotsky, J. Bielefeld, L. Huang, M.N. Baliki, G.D. Iannetti, A.V. Apkarian

## Abstract

High-profile studies claim to assess mental states across individuals using multi-voxel decoders of brain activity. The fixed, fine-grained, multi-voxel patterns in these “optimized” decoders are purportedly necessary for discriminating between, and accurately identifying, mental states. Here, we present compelling evidence that the efficacy of these decoders is overstated. Across a variety of tasks, decoder patterns were not necessary. Not only were “optimized decoders” spatially imprecise and 90% redundant, but they also performed similarly to simpler decoders, built from average brain activity. We distinguish decoder performance when used for discriminating between, in contrast to identifying, mental states, and show even when discrimination performance is strong, identification can be poor. Using similarity rules, we derived novel and intuitive discriminability metrics that capture 95% and 68% of discrimination performance within- and across-subjects, respectively. These findings demonstrate that current across-subject decoders remain inadequate for real-life decision making.

## INTRODUCTION

A whole body of neuroimaging literature—largely published in highly influential journals—either explicitly claim, or strongly imply, that thinking is no longer private. By “optimizing” functional magnetic resonance imaging (fMRI) brain scan results, these studies profess to universally decode mental states: feelings, thoughts, decisions, intentions, and behaviors ^1-3^. Thus, neuroscience seems to have broken the code of mental states, in turn proclaiming the ability to “read the brain” of every human being. Here, we systematically examine the validity of such claims.

Decodability—how discernable a mental state is, given a brain activity pattern—is predicated both on the brain activity properties of the task being discerned as well as the goal of the decoding. Intuitively, decodability is analogous to discerning a breed of dog; breeds that look more similar will be harder to distinguish. The literature claims “optimized” decoders can (1) discriminate between mental states, (2) identify mental states, and (3) capture additional state-related measures (stimulus or perception intensities). These goals can be more tangibly elucidated through the dog breed metaphor: Consider a pug (a *decodee*) and a French Bulldog (a *comparator*)—two breeds that may look alike. If one is familiar with the unique physical features of a pug—small stature, short snout, wrinkled face, folded ears, curled tail, etc.—then such features can serve as the *decoder* for a pug. This decoder can then be used to perform the three decoding tasks. Specifically, discrimination is akin to deciding which dog is a pug when the pug and French Bulldog are next to one another. Conversely, identification is akin to saying whether a single dog is a pug when there are no other dogs around. On the other hand, capturing a continuous measure, such as perceived intensity of a state, is much like trying to judge a dog’s age. While discrimination and capturing continuous measures have been discussed and illustrated for various mental states, less attention has been given to identify a certain mental state from a given pattern of brain activity.

The pattern of mental state decoders arises from weights assigned to its constituent voxels; for this reason, we call them *fixed-weight decoders*. Voxel weights are derived in three stages. First, general linear models (GLM) generate a brain *activity map* (correlation between the activity in each voxel and the task). Second, GLM is used to contrast the activity maps from a task or state of interest (a *decodee*; e.g., pain) to one of no interest (a *comparator*; e.g., touch), and its results are thresholded (a *contrast map*). Finally, “machine learning” models are used to tune the weights in the contrast map to optimize its performance ^4,5^; the result is a relatively sparse, fixed-weight decoder with a fine-grained pattern (an *“optimized” decoder*). It is tacitly assumed that each stage improves performance of the decoder by uncovering better distributed patterns of neural ensembles related to the mental state, and as a result, detailed spatial patterns confer predictive value, as explicitly stated, “the pattern of activation, rather than the overall level of activation of a region, is the critical agent of discrimination” ^5^. This concept is now expounded for diverse topics across many labs ^3,5-11^.

The concept that across-subject “optimized” decoders are able to capture mental states across different individuals violates basic neuroscientific principles. The technical and biological requirements of such decoders are quixotic, as they imply the existence of a fixed, exclusive, universal brain activation pattern for each and every mental state—a one-to-one correspondence between subjectivity and objective brain patterns. Such invariant brain-to-mind models imply a common neuronal firing pattern across billions of neurons, which is unique for every mental state and shared across all humans. This invariance is purported in spite of large inter-subject variability in gross brain anatomy, as well as of differences in genetics, lifestyles, lifetime experiences, and associated memory traces ^12,13^; all of which would carve the individualized brain activity of subjectivity (for a discussion on the topic from the viewpoint of fMRI analysis, see ^14^). If a trivial, fixed relationship exists between subjectivity and brain activity, such “optimized” decoders also raise strong ethical and legal concerns regarding their ability to invade mental privacy ^15^, and also would be incongruent with commonly accepted philosophical constructs of subjectivity ^16^.

Our principal aim was to evaluate the performance and necessity of “optimized” decoders relative to more parsimonious approaches (e.g., GLM maps). After rigorously evaluating the performance of “optimized” decoders, we sought to understand fixed-weight decoders from a more general perspective: What determines and constrains decodability?

## RESULTS

### Overview

Our investigation began with three published pain decoders. Both qualitatively and quantitatively, these decoders were markedly different from one another (**Fig 1**). Despite these differences, on average, their ability to discriminate pain from non-pain states, across four published studies (*N*=113) ^4,5,8,17^, was nearly identical. To understand how disparate decoders could perform similarly, we parametrically perturbed each of the decoders and tested its performance. Perturbations consisted of 1) searching for brain locations privileged for decoding pain; 2) randomly using subsets of voxels from select regions; 3) using subsets of voxels based on their weights; and 4) spatially smoothing (and thereby modifying voxel weights). The analysis demonstrated that the tested decoders were ∼90% redundant in space and, remarkably, that their weights were superfluous for successful discrimination (**Fig 2**). Similar results were obtained for stimulus-perception mapping (**Fig 3**). Overall, we observed that sparse, location-only based decoders were sufficient for discriminating pain.

**Figure 1.**
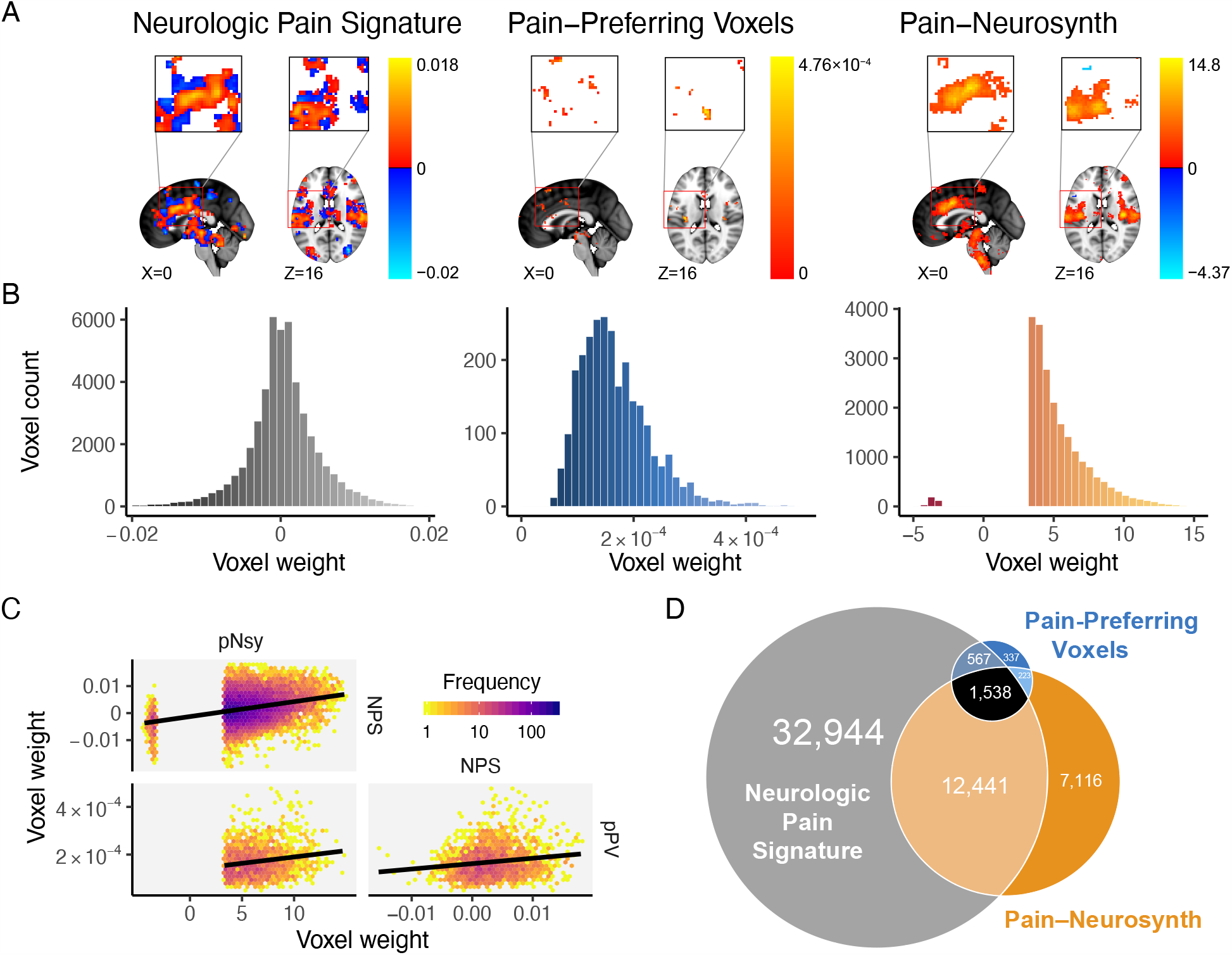
Spatial properties for three decoders, which are supposed to distinguish pain from other mental states, are distinct from each other. **(A)** Location and voxel-wise weight patterns of the three pain decoders (respectively abbreviated NPS, pPV, and pNsy). **(B)** Weight distributions of all three decoders are distinct. NPS weight values are distributed around zero; pPV has no negative weights; pNsy has only a few negative weights. **(C)** Pairwise correlations between weights of the three decoders. Lines depict total least squares regression fits. All three correlations are weak (*r*_NPS-pPV_ = 0.16; *r*_pNsy-NPS_ = 0.30; *r*_pNsy-pPV_ = 0.18). **(D)** Euler diagram depicts relative size of each of, and spatial overlap between, the three decoders.

**Figure 2.**
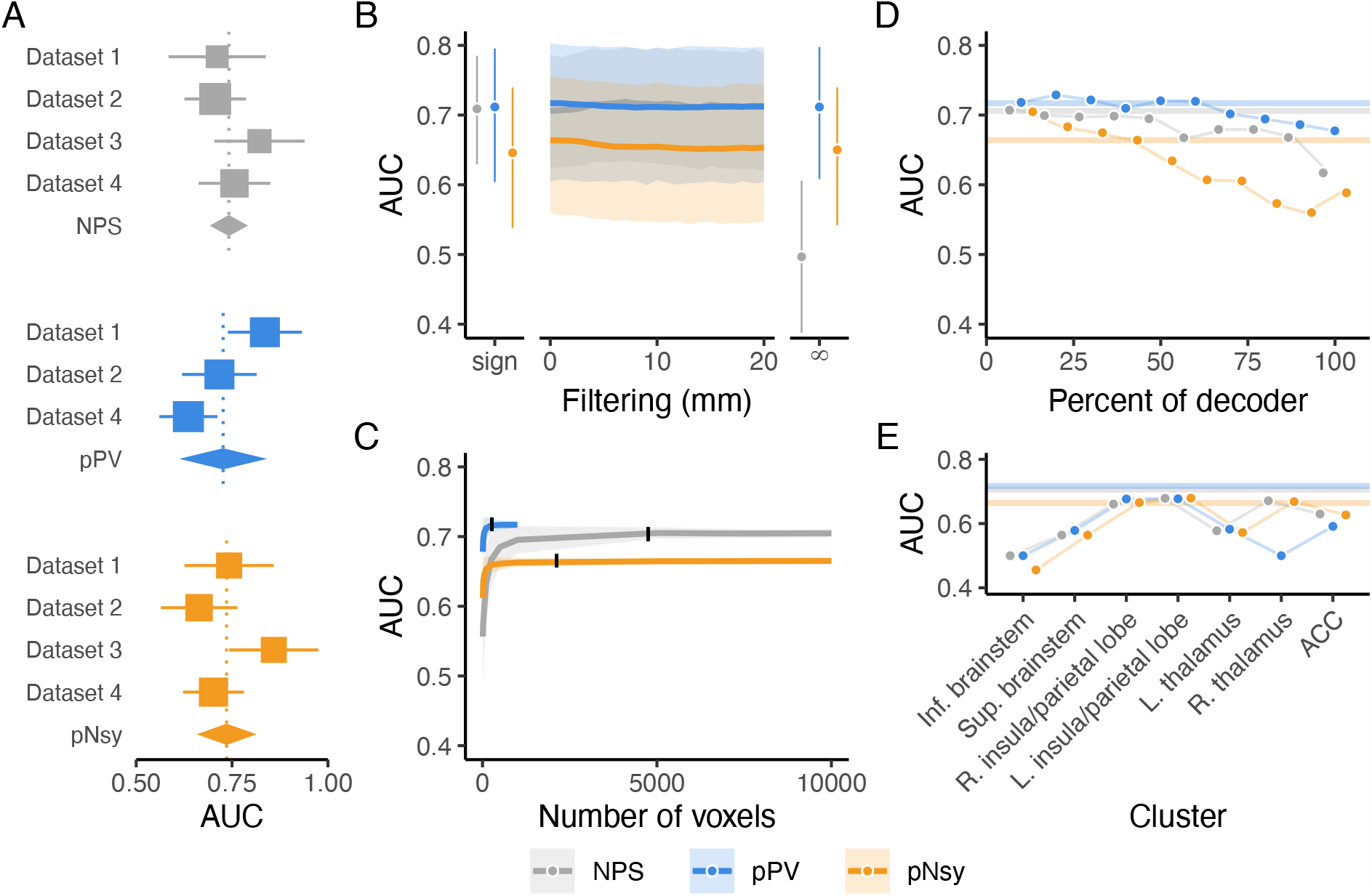
Discrimination performance is similar for all three pain decoders and is a function of voxel locations, not weighted patterns. **(A)** Meta-analysis of across-subject discrimination performance (AUC, chance = 0.5) for decoding pain from non-pain mental states for each of the three decoders, tested only for datasets independent of decoder derivation. On average, all decoders perform similarly. Square sizes indicate meta-analytic weight. **(B-C)** Across-subject decoding of pain from touch. **(B)** performance does not change when decoder pattern weights are distorted with increasing-size spatial smoothing. Sign = sign of voxel weights with 0 filtering, rendering decoder voxel values to only 0, −1, +1; filtering σ= 0–20 mm; ∞ = location-only. **(C)** Decoder performance depends only on a very small number of voxels, indicating information redundancy. The number of voxels constituting each decoder was systematically increased (from 10 voxels to the full decoder) and performance assessed for random samples of each size. 10% of each full decoder’s voxel count (black ticks) discriminates pain from touch equivalently to the full decoders. Shades are standard deviations for spatial uncertainty, ignoring across-subject uncertainty. **(D)** Decoders were constructed using 10% of the voxels in the full decoders, with voxels selected in order of their absolute magnitude (see **Fig S7**). The voxels with the highest absolute weights do not necessarily discriminate better than voxels with lower magnitudes, with the exception of pNsy in this dataset. **(E)** Selecting voxels based on their anatomical locations revealed that single regions (e.g., L. insula/parietal lobe) can discriminate similarly to the full decoders. Bars and shades are the 95% confidence intervals [CI] of means, except in **C**, where shades indicate standard deviations associated with permutation variability. In **D** and **E**, colored bars indicate the AUC of the full decoders.

**Figure 3.**
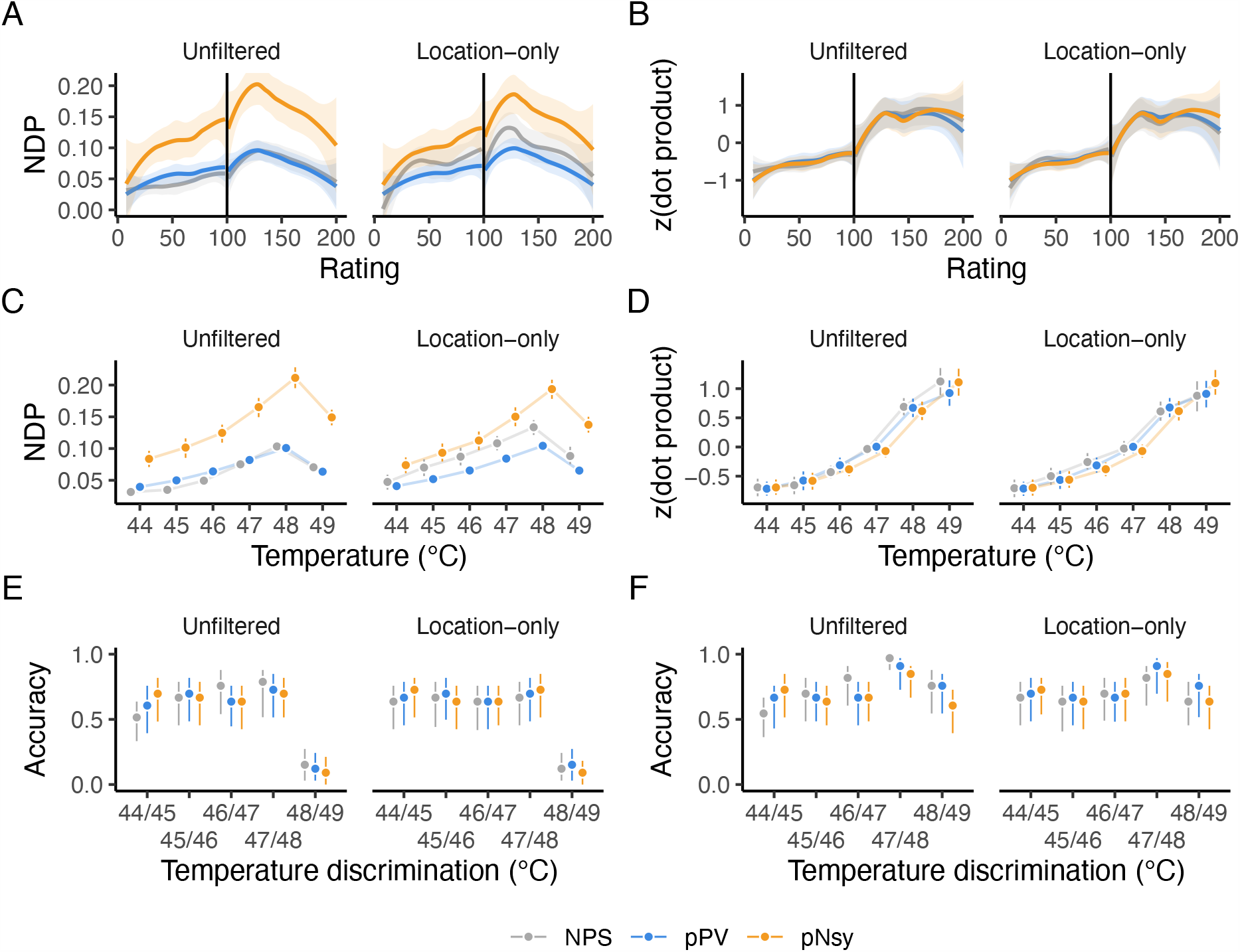
All decoders perform stimulus-perception mapping similarly, both with and without voxel weights. All three pain decoders perform equivalently, when location-only decoders are compared to the unfiltered-decoders, in mapping pain and heat perception ratings **(A–B)**, mapping painful stimuli **(C–D)**, and discriminating between pairs of painful stimuli **(E–F)**. Nonmonotonic relationships indicate that the decoders cannot reliably predict subjective ratings or stimulus intensity. Vertical lines in **A** and **B** indicate the transition from heat (< 100) to pain (> 100). The dot products in **B, D**, and **F** were *z*-scored within each decoder for presentation purposes.

To further generalize this finding, we examined decoding properties for cognitive domains other than pain, where dedicated brain tissue is better established; namely, a reading task and a listening task (two publicly available datasets, *n*=14 and *n*=213 subjects, respectively) ^18,19^. We compared decoding performance between GLM and “optimized” decoders, before and after constraining to location-only. Our results closely resembled those for decoding pain (**Fig 4**).

**Figure 4.**
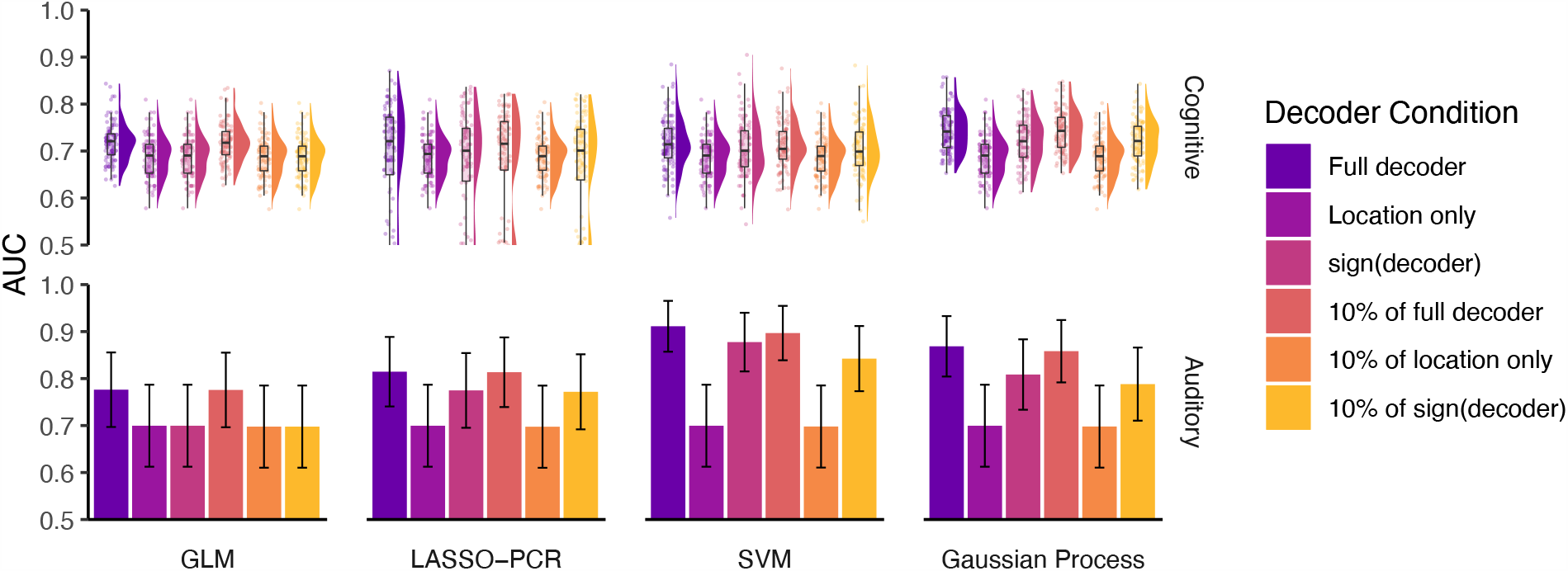
Different implementations of cognitive and auditory decoders perform similarly regarding discrimination performance and are robust to perturbations. We constructed decoders using general linear modeling (GLM), least absolute shrinkage and selection operating with principal components regression (LASSO-PCR), support vector machines (SVM), and Gaussian processes to decode **(top)** cognitive ^19^ and **(bottom)** auditory tasks ^18^. Much like the pain decoders, these decoders performed similarly and better than chance (chance = 0.5 in both), and were relatively robust to perturbations. Just 10% of each decoder was enough to capture its full performance, and even extreme perturbations, such as 10% of the binary decoder or 10% of signed decoder, had little effect on performance. Error bars are the 95% confidence intervals of the AUCs. *Nota bene*, in the auditory task, discrimination performance is better with SVM and Gaussian Process than with GLM or LASSO-PCR. We suspect these differences are a consequence of specific instantiations of overfitting. We observed similar decoder-dependent performance variations for the pain decoders as well (see **Fig 2A**); yet, in further analyses none showed superiority over the others. In the auditory task, and for both SVM and Gaussian Process decoders, we also observed appreciable performance decrement for location-only and for 10% location-only decoders. This too was observed in the pain decoders. Like with the pain decoders, here, we also observed that sign-only decoders and 10% sign-only decoders performed similarly to the full decoders, again suggesting that negative weights at large scales can influence decoder performance.

The brain imaging literature commonly accepts that if a decoder can adequately discriminate between a decodee and a comparator, then it is also useful for identifying the mental state associated with the decodee. We tested this concept for both pain and listening tasks. Despite discrimination being possible and robust to perturbations, all decoders performed poorly and relatively similarly when trying to *identify* the decodee mental state (**Fig 5**).

**Figure 5.**
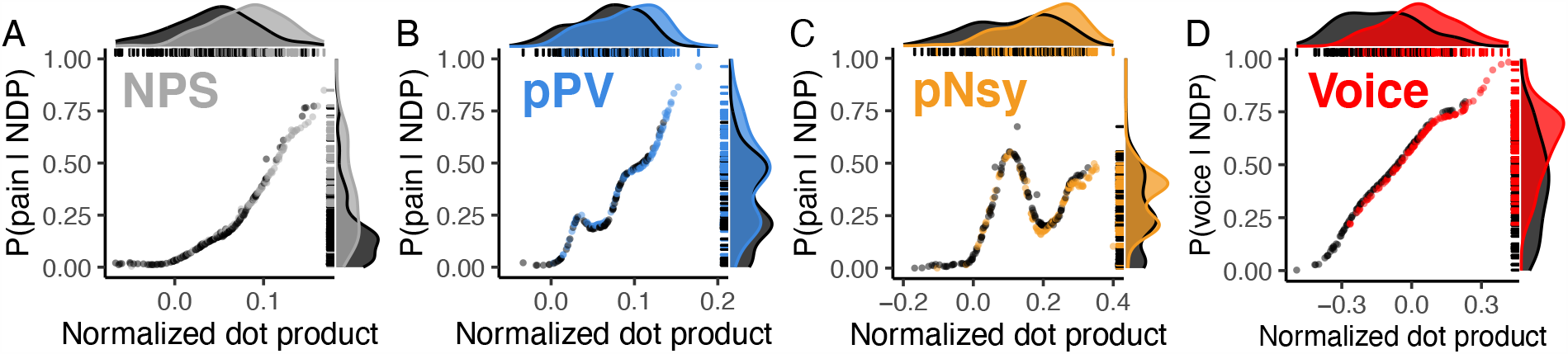
Identification of mental states shows poor predictability. Three pain decoders (NPS, pPV, and pNsy in **A–C**) and a voice decoder (**D**) were used to test identification for mental states. *x*-axes are the normalized dot products between decoder and decodee, while *y*-axes are the posterior probability of being in pain (**A–C**) or listening to voices (**D**). Distributions of normalized dot products and posterior probabilities include both the decodee (light grey & colors) and comparator (dark grey) tasks. (**A–C**) Normalized dot products of the pain condition span the entire distribution of comparator normalized dot products, and as a result, pain is not strongly isolated from the comparator conditions. Quantitatively, this is evidenced by the strong decodee-comparator overlap for (**A**) NPS (overlap (95%CI) = 68% (59–82)), (**B**) pPV (79% (73– 90)), and (**C**) pNsy (73% (66–84)). This is reflected in the Bayesian model, which shows similar probabilities of being in pain for both pain and pain-free conditions (each dot/line). To this end, all three decoders perform similarly, and cannot unequivocally *identify* pain, as indicated by their low sensitivity/specificity (when specificity/sensitivity=0.95) of (NPS, **A**) 0.19/0.25, (pPV, **B**) 0.25/0.17, and (pNsy, **C**) 0.17/0.27. (**D**) In contrast to pain, a contrast map decoder for identifying when a participant is listening to human voices separates more clearly the normalized dot products of the decodee (red) from comparator (dark grey), but still performs poorly (overlap = 54% (46–66)). This separation is reflected in the Bayesian model, which shows high probabilities when individuals are listening to human voices and lower probabilities when they are not. To identify the mental state of listening to voices based on NDP with a specificity of 0.95, one would have a sensitivity of 0.19. Conversely, to identify the same mental state with a sensitivity of 0.95, one would have a specificity of 0.48.

The results of our perturbation analyses led us to explore the limits of decoding. If perturbed and simplified decoders can perform similarly to the original “optimized” decoders, can we further simplify decoders and also quantitate decodability? To address the former question, we built pain decoders using GLM maps for noxious stimuli. These GLM decoders performed similarly to “optimized” decoders, with within-study performance being slightly superior to across-study performance (**Fig 6a,b**). We extended these findings to quantify within-and across-subject decoding using four different tasks, repeated up to 12 times per subject in 14 subjects ^19^. This study design provides the opportunity to calculate discriminability as a function of similarity measures from the decoder, decodee, and comparator, for both within- and across-subject decoding. Although performance was not consistently better for within-subject discrimination, variation in performance could be largely explained by within-task homogeneity and between-task heterogeneity, allowing us to propose decoding rules (**Fig 6c,d**), which worked better for explaining within-compared to between-subject discriminability. These results present convergent evidence that 1) discrimination decoding is limited by GLM results, where sparse location-only maps contain sufficient information; 2) identification is harder than discrimination; 3) similarity measures almost fully account for the variance of within-subject discrimination decodability, which degrades in across-subject discrimination.

**Figure 6.**
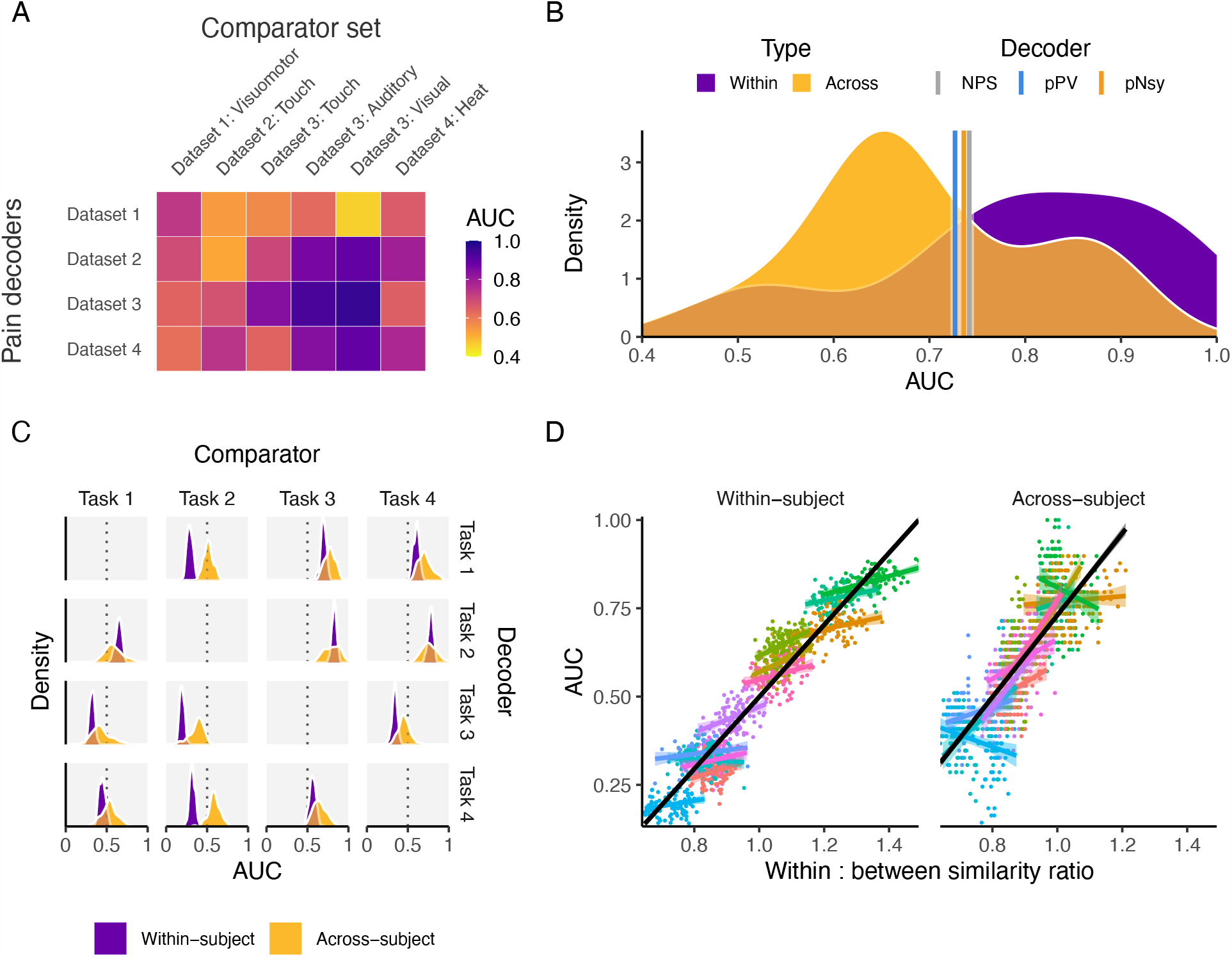
Decoders constructed from activity maps perform similarly to pattern-based decoders, and are dependent on both decodee and comparator properties. **(A)** Performance of four activity map decoders, based on the across-subject averaging for pain tasks, to differentiate pain from six other mental states. **(B)** Among the activity map decoders, within study performance is slightly higher but extensively overlaps with across study performance. Meta-analytic estimates of performance for NPS, pPV, and pNsy are within 0.4 standard deviation from the average performance of both within and across study activity map decoders. **(C-D)** Properties of activity map decoders are examined within and across subjects as a function of a cognitive task ^19^. **(C)** Decoders (rows) are built from four cognitive tasks, tested on remaining three (columns), in a within subject and across subject design. Within subject performance is always more consistent (i.e. it has smaller variance) but not necessarily greater than across subject. For example, the within subject performance is always superior to across subject when using task 2 as the decoder. The inverse is true when task 2 is the comparator, implying strong task dependence. **(D)** Decoder performance linearly scales with the ratio of decodee similarity to decodee-comparator similarity (based on normalized dot product), for within- and across-subject comparisons. Because discriminability depends on this ratio of similarities, they can be viewed as rules for decoding. Each color in (**D**) represents a decodee-comparator pair of tasks 1-4 in (**C**); each point is a permuted sample; each colored line is the regression within a decodee-comparator pair; and the black line is the regression across decodee-comparator pairs.

### Exploring Established Decoders

We started by assessing “optimized” decoders for pain using the Neurologic Pain Signature (NPS, constructed using LASSO-PCR ^5^) and Pain-Preferring Voxels (pPV, constructed using SVM ^4^). In addition, we used a meta-contrast map as an alternative decoder, Pain-Neurosynth (pNsy), which is the meta-analytic association test result for the term “pain”. This contrast map is based on 516 studies containing the word “pain” in the abstract, and contrasting them with the remaining 13,855 neuroimaging studies (using the public tool Neurosynth ^20^; see Supplementary Methods). We first compared the spatial and weight properties of these three decoders. Although all decoders cover approximately the same brain regions (**Fig 1A**), the distributions of their voxel weights are distinct (**Fig S1**), the numbers of their constituent voxels are vastly different, their pairwise correlations are weak, and their spatial overlaps of voxels are relatively small (**Fig 1, B-D**).

### Discrimination Performance for Pain is Similar Between Diverse Decoders

To enable decoding, we assessed the similarity between the decoder and decodee or comparator using the normalized dot product (NDP; +1 indicates total/maximal pattern similarity, 0 indicates orthogonal patterns, −1 indicates anti-similarity). Discrimination was assessed using the area under the receiver operating characteristic curve (AUC) from the two distributions of NDPs. AUC is an indicator of discriminability since it can be interpreted as the probability of a randomly sampled decodee NDP being greater than a randomly sampled comparator NDP, implying a direct comparison. Conversely, identifiability was assessed using distributional overlap, with greater overlaps indicating poorer identifiability. Points contained within the area of overlap are equally likely to be in the decodee and comparator distributions, and thus, are not identifiable. Together, these metrics served as the basis for decoding performance throughout this study.

We examined the performance of the three pain decoders for discriminating pain states from control tasks, and for capturing stimulus/perception properties, in 4 published studies from 3 labs (*N*=113 subjects). Despite marked spatial and weight distribution differences, average discrimination performance (meta-analysis, AUCs pooled across datasets and various comparators) were approximately equivalent (AUC ≈ 0.73 for all three; **Fig 2A**). This equivalence is remarkable and informative: it implies that very different models may nonetheless yield similar average performance, suggesting that their detailed properties do not constrain decodability. Notwithstanding similar average performance, the decoders performed differently for particular datasets, indicating that decoding performance has a specificity component which can likely be explained by brain region-specific dependences.

### Pain Decoders Are Robust to Spatial Perturbations

#### Search for Brain Locations Privileged for Decoding Pain

To test whether there are privileged locations for decoding pain, we created clusters of voxels using the common space across the three pain decoders and tested discrimination performance for all three decoders across the four tasks (**Fig S2**). For any given study, multiple clusters from multiple decoders performed equally well and matched the performance of the full decoder. This result suggests that no single cluster was consistently more specific for decoding pain than other clusters.

#### Spatial Smoothing and Voxel Weights

To investigate whether discrimination performance relies on the *fixed-weight* nature of the voxel patterns, we measured performance when these patterns were degraded (1) by spatial smoothing and (2) by discarding their weights. Remarkably, decoding performance was minimally affected by either procedure (**Fig 2B, Figs. S3-S6**). This result clearly demonstrates that the fine-grained pattern of weights in “optimal” decoders added no value to performance (with a few exceptions, **Fig S3**); rather, voxel locations alone were sufficient for discrimination.

#### Number of Voxels

Given the three decoders’ structural differences and yet similar performances, it can be virtually ruled out that all voxels composing these decoders are necessary to achieve satisfactory discrimination. To characterize the minimum number of voxels necessary to decode the pain state, we created sets of new decoders by randomly selecting subsets of voxels from each decoder. Surprisingly, we attained the original decoding performance when only using a random 10% of the total number of each decoder’s constituent voxels (**Fig 2C**). We replicated this finding on all datasets and for all three decoders, both in their original form and when only using their signed voxel locations (**Figs. S4–S6**). We further explored the relationship between voxel weights and performance by first binning voxels by their absolute weights and then constructing a set of decoders using the voxels in each bin (see **Fig S7**). Again, we observed only a minimal degradation of performance with decreasing voxel weights for all decoder-dataset combinations (with some degradation seen mainly for unfiltered NPS at highest binning, **Figs. S8–S9**). Together, these results demonstrate that the information within the set of voxels present in the decoder, decodee, and relative to the comparators was highly redundant and essentially independent of the decoders’ weights.

#### Fixed, Multi-voxel Patterns Confer No Added Value to Stimulus/Perception Intensity Decoding

Wager, et al. ^5^ used an “optimized” decoder, NPS, not only to discern the dichotomous presence of pain, but also to claim that NPS can capture stimulus intensity and perceptual ratings from brain activity. To this end, we tested the ability of the three pain decoders to capture stimulus and perception properties. We used data from a study where nonpainful and painful stimulus, perceptual responses, and their associated brain activity were available ^5^. All three decoders (NPS, pPV, and pNsy), whether unfiltered or infinitely filtered (location-only), performed similarly for capturing perceived pain ratings (**Fig 3A,B**), for reflecting the intensity of the thermal stimulus (**Fig 3C,D**), and for discriminating between pairs of painful stimuli (**Fig 3E,F**). The similar performance between unfiltered and location-only decoders demonstrates that response trends arise as a consequence of the intensity changes of the decodee rather than the weight distribution of the decoder. Moreover, the discordant performance between NDP (nonmonotonic, **Fig 3A, C, and E)** and dot product (almost monotonic, **Fig 3B, D, and F)** suggests that previously reported results ^5^ were primarily due to an increase in the magnitude of brain activity in specific regions, but in a way that becomes less similar to the decoder. Yet, both NDP and dot product were robust to the removal of voxel weights. These results again refute superiority of “optimal” decoders above that of a meta-contrast map decoder, which was derived from GLM results. This reinforces the notion that location-only performs sufficiently, and that useful information is provided only by the decodee activity within the locations where a decoder has non-zero weights.

### Cognitive and Auditory Decoders Are Similarly Highly Redundant

So far, we have shown that popular “optimized” pain decoders, as well as a meta-contrast map used as a decoder, are able to maintain their full performance after being perturbed and degraded, indicating that much of the information contained within them is superfluous. One worries that the findings may be specific to the modality studied, as pain and nociception are sensory systems for which no dedicated neocortical tissue has been uncovered in the cortex ^21^. As a result, there is long-standing debate as to specific or distributed encoding of pain perception (e.g., ^22^; cf. ^23^). To broaden our findings, we examined whether the uncovered principles apply to decoding for audition and language. Primary and secondary auditory cortex ^24,25^ are in close proximity to the somatosensory regions examined above for pain, while language representation with dedicated and functionally specific tissue is unique to humans ^26^. We used data for language ^19^ and auditory ^18^ studies to construct decoders using task-specific contrast maps, SVM, LASSO-PCR, and Gaussian processes (our contrast maps closely resemble those reported in the original studies, **Fig S10– S11**; see Supplementary Methods). Our findings are entirely concordant with those for the pain decoders, in that all of the constructed decoders show similar performance, which was maintained after extreme perturbations (e.g., sign or location-only) (**Fig 4**), with only a few exceptions (see **Fig 4** comments). These findings generalize and provide compelling support for our main result: “optimized” decoders are highly redundant, and decoding primarily exploits information contained within voxel locations, independent of voxel weights. Moreover, task-specific GLM contrast maps are sufficient, implying that the meta-contrast maps are also not necessary.

### Identification Remains A Challenge

The ability of “optimized” decoders to *identify* mental states is repeatedly asserted in the literature ^3,5-11^, but to our knowledge, remains untested. Specifically, the sensitivity and specificity reported in previous works are estimated by changing decoder response thresholds for different pain stimuli ^5^. If “optimized” decoders are used with the objective of identification, then they should be able to pinpoint the specific mental state solely from the similarity between the decoder and decodee, and, crucially, in the absence of a comparator and without such a threshold tuning. In other words, identification should be based on a single observation and what we (or the decoder) “know(s)” about the world. Therefore, instead of AUC, which implies a comparator, we tested identifiability by calculating distributional overlap between the states of interest and no interest. Distributional overlap provides the range of equal probability of belonging to the state of interest and state or states of no interest; here, equiprobability implies unidentifiability. In addition, we were interested in assessing performance at the individual level. To do so, we calculated the probability of a subject being in a specific mental state given that subject’s brain activity map. We thus calculated distributional overlaps and state probabilities to assess the ability of decoders to identify mental states.

Identification of pain states was similarly poor across the three pain decoders explored: overlaps between states of interest and states of no interest were high (≥ 68%) and the probabilities of being in pain (when actually in pain) were low (median posterior probability ≤ 0.5) **(Fig 5a–c)**. These results paint a markedly different picture than the discrimination results, which simply show that NDPs *tend* to be greater when individuals are in pain; evidently, adequate discrimination does not translate to identification. We built upon these findings by using the task-specific contrast map decoder to decode audition of vocal versus non-vocal sounds ^18^. While the performance of the voice decoder was better than that of the pain decoders (overlap = 54%), it was still inadequate, as over half of the data was unidentifiable (**Fig 5d**). The slight superiority of the voice decoder relative to the pain decoders may have several explanations, including the homogeneity of the training and test sets used for the voice data, or simply that some tasks are easier to identify than others. In any case, regardless of the mental state tested, identification remained unreliable and thus is currently not feasible with fixed-weight decoders.

### Brain Activity Maps Are Sufficient for Discrimination

The similarity in performance achieved by meta-contrast maps or task-specific contrast maps and “optimized” decoders prompted us to take another step back in the decoding derivation process. Given that pNsy is a composite of contrasts from many studies (i.e., a meta-contrast pain decoder) and its decoding performance was similar to “optimized” decoders (NPS and pPV), we assessed whether an *even* simpler construct—pain activity maps—was sufficient to decode the state of being in pain. In other words, if no performance is lost by using contrast maps, would task-derived GLM maps suffice as simpler but adequate decoders? Brain activity map decoders were created using the average brain activity for each study’s pain task. Each activity map decoder was then used to discriminate pain using the left-out brain activity maps of subjects both within and between studies (**Fig 6A**). Remarkably, these decoders performed comparably to the ones presented hitherto (NPS, pPV, and pNsy), with an average within-study AUC of 0.79 and between-study AUC of 0.69 (cf. ∼0.73 for the fixed-weight decoders; **Fig 6B**). The lack of clear superiority of “optimized” decoders relative to a meta-contrast map, and even simple activity map decoders, casts serious doubt on the predictive and epistemological value of the more complex “optimized” decoders. These results also raise the salient question: If decoding can be approached in so many different ways, what actually determines decodability?

While decoding is difficult, decodability itself is likely predictable, yet to our knowledge remains unexplored. To build upon our breed metaphor, some dogs exhibit features that largely overlap with other dogs, such as the stature, color, and flat-faced features of pugs and French Bulldogs. Similarly, the mental state of “being in pain” shares many features with other states; for example, unpleasantness, behavioral relevance, and saliency ^27^. Therefore, the primary challenge of decoding is to tease apart these overlapping features. For this reason, it seems logical that the similarity of activity maps within and between the decoder, decodee, and comparator would determine decoding performance. If the decoder is built from activity maps that are dissimilar, the resulting average map would have a low signal-to-noise ratio; if the decodees or comparators are dissimilar, then we can expect a greater variance in NDPs; and if the decodees and comparators are similar to one another, then they will have a lot of overlap and be difficult to tease apart. This logic implicates the neuroanatomical and physiological assumptions previously mentioned, as heterogeneity across individuals should decrease similarity, making the NDPs more variable and thus more difficult to discern. Using similarity metrics that reflect these relationships, we attempted to explain decodability.

Until now, we have primarily focused on decoding across-rather than within-subjects. Intuitively, it is apparent that, for many of the reasons elaborated above, decoding mental states should be more successful within-subjects compared to across-subjects, as has been formulated by others ^28,29^. However, no systematic analysis of this notion has been performed using fixed-weight decoders. Therefore, we investigated this question using data well-suited for the question: fMRI data collected from 14 subjects who completed four cognitive tasks, each with 12 replicates ^19^. These repetitions enabled the comparison of decoder performance within- and across-subjects. As expected, decoding performance is more precise (smaller variance) within-subject (**Fig 6C**), but, interestingly, not necessarily better (greater average AUC). We investigated whether the ratio of decodee to decodee-comparator similarity (or within:between) can be a possible natural metric of why some decoders are more efficacious than others. Higher performing decoders showed greater within:between ratios than lower performing decoders (**Fig 6D**). Similarly, decoder similarity—the average NDP of all pairwise combinations of a decoder’s constituent activity maps, a measure of reliability—could also explain much of the decoder performance, and in support of our previous conclusions, this relationship is largely unaffected by thresholding and binarizing the decoder (**Fig S12**). Further exploration showed that decodability, especially within-subject, is strongly predicated on these similarity metrics (**Fig S13–S14; Table S1**). Decodee similarity, together with decodee-comparator similarity, is strongly predictive of discriminability, accounting for up to 95% of the variance in AUCs. Our similarity metrics almost entirely explain within-subject decodability, but only about 68% of AUC variance in across-subject decoding. This result may speak to the assumptions violated by across-subject decoders, in that a similarity score across-subjects is less interpretable than one calculated within a single subject since variance (e.g., brain anatomy) may be converted to bias (making all brains fit the same template) during image preprocessing and registration.

## Discussion

In this study, we asked what the determinants, and limits, of decoding mental states are. We primarily emphasized decoding pain, as this is the modality where the most emphatic claims have been made and where the “optimized” decoders seem to have become accepted as enabling “mind-reading” ^3^. For pain, audition, and language tasks, the locations of a small subset of GLM-derived voxels were sufficient for achieving a discrimination of AUC ≈ 75%, and a long list of machine learning tools could not consistently improve upon this performance. We also showed that, in contrast to discriminating between states, identification of a given perceptual state is much harder. For the first time, we advanced the concept of quantifying discriminability using a simple similarity metric, the NDP, with which we provide models for within- and across-subject discrimination. The latter analyses indicated that discriminability depends not only on the decoder, but also on similarity between the decodee and comparator. Finally, we showed that, even in an example where within-subject discrimination was almost fully modeled with similarity properties, there was a considerable decrease in the variance of across-subject discrimination that could be explained. In doing so, we established limits of decodability based on the most popular linear models currently used in fMRI literature.

Limitations of across-subject decoding and reverse inference have been acknowledged by others. For example, the latest evidence shows that brain-behavioral phenotype associations seem to become reproducible only with sample sizes of *N* ⪆ 2,000 ^30^. Yet, the extent of these limitations has not previously quantified, nor has decodability been modeled. Multiple approaches have been initiated to overcome these limitations. The simplest is to constrain functional studies to within-subject investigations, thus bypassing the idiosyncrasies of anatomically aligned, group-averaged results, but this approach also obviates across-subject decoding. The approach has been used in various topics, including subject-specific localizers in vision ^31^ and language studies ^32^.

Perhaps the most widespread method is the multivoxel pattern analysis (MVPA). MVPA looks for statistical evidence for *information* contained within groups of voxels (functional/anatomical regions of interest or searchlights) ^33^. MVPA has been successful in decoding diverse brain states from fMRI activity patterns; for example ^4,34-40^. MVPA typically uses subject-specific classifier models, and as a result, its accuracy drops when predicting other subjects’ responses ^28,29,38^. To extend MVPA results to across-subject applications, and to improve on anatomical alignment, Haxby and colleagues ^33,41,42^ developed an across-subject, high-dimensional, functional alignment technique, named hyperalignment. It has previously been shown that hyperalignment, coupled with MVPA, improves across-subject response classification to levels that are comparable to, or even better than, those seen for within-subjects ^29,38^. To do so, hyperalignment exploits the temporal variability of stimulus-evoked brain activity, yet it is designed to enable alignments based on diverse brain signals ^29,33,39^. Therefore, we explored whether hyperaligning GLM brain activity maps would enhance across-subject decodability. In contrast to previous work, our preliminary results did not show improvement between hyperaligned and anatomically aligned across-subject decodability (data not shown). Still, it is possible that variants of hyperalignment may be useful in brain activity-based across-subject decodability (e.g., ^43^).

Our principal finding is consistent with the MVPA literature. A recent across-subject study used MVPA (without hyperalignment) to uncover circuitry associated with pain relief commonly seen for eight different types of analgesics ^44^. Their study identified brain locations involved in analgesia for multiple drugs, and thus it is consistent with our main conclusion: location and not fixed patterns are sufficient decoders. In fact, in general, MVPA identifies within- or across-subject brain locations, at macro- (voxel level) or micro-scale (sub-voxel level, ^45^), where a certain discrimination or calculation is possible. The level of discrimination will ultimately be constrained by differences in functional-anatomical coupling across individuals, in turn leading to distinct results within- and across-subjects ^46^. Thus, specific underlying patterns may not be identified, which again is consistent and complementary to our main conclusion.

Beyond promoting reverse inference, fixed-pattern decoders, also labeled as “signatures” or “neuromarkers” ^2,3^, are purported to 1) unravel neural encoding of psychological constructs, 2) improve decodability, 3) enable validation across studies and labs, and 4) provide falsifiable models. Since such decoders do not outperform brain activity map-based decoders, we contend that the aforementioned assertions are untenable. It follows that fixed-pattern decoders do not provide a defensible path forward for constructing a brain activity-derived ontology of mental constructs ^47^.

Our demonstration that overlaying linear machine learning optimization algorithms did not improve on linear contrast-derived decoders is not surprising. Indeed, similar conclusions have been reached in other domains, such as medicine ^48,49^. Moreover, our findings support the idea that neuroimaging has not saturated the performance of simple linear models ^50^. The reasons for this are manifold, and from a modeling viewpoint, it has been argued that the added value of linear “machine learning” techniques is often small, exaggerated, and does not translate into practical advantages ^51^. Although unsurprising given the aforementioned work in this area, the apparent stark discrepancy between our findings and those in the literature warrants explicit explanation.

How do we explain the discrepancy between our results and the literature, even when the same decoder is used on the same data ^5^? We cannot escape the conclusion that “optimized” decoders are superfluous models. Indeed, Wager and colleagues have also observed similar performance across several pain decoders, including NPS and pNsy ^5,52^. Moreover, the use of arbitrary performance metrics (here we base all analysis on NDP), lack of a control (comparison to GLM modeling), and commonly mixing within- and across-subject performance metrics all seem to mislead and propagate grandiose assertions ^53^. In stark contrast, here we show that linear models—contrast and activity maps—are capable of maximizing prediction, while being readily available and maintaining interpretability. Yet, across-subject decodability remains complex; only brain location adds value, and depends on within and between similarity of decoder, decodee, and comparator. These findings advance the general principles of decoding mental states. Importantly, the limited and inadequate performance of fixed-weight across-subject decoders, especially regarding identification, pose strict bounds on their utility in the domains of medical and legal decision-making.

## Supporting information

Supplemental Results

## Acknowledgments

We would like to thank Dr. Thorsten Kahnt and Apkarian lab members for providing their thoughtful feedback.

## Authorship Contributions

RJ, ADV, MNB, GDI, and AVA conceived the idea; RJ, ADV, JB, and LH performed the analyses; RJ, ADV, and AVA drafted the manuscript; RJ, ADV, JB, GDI, and AVA edited the manuscript; and all authors approved the final manuscript.

