## Supplemental Methods for "The Hard Limits of Decoding Mental States: The Decodability of fMRI"

### Materials and Methods

Datasets

6 datasets were used in this paper; all are part of published studies, and were either provided by their authors (Datasets 1, 2, 3, 4, and 5) or downloaded from public repositories (Dataset 6). Datasets 1, 2, 3, 4, and 5 consist of voxel-wise, whole brain, task dependent GLM analysis activation maps (beta maps). Dataset 6 consist of BOLD timeseries, and were processed using standard fMRI pre- and post-processing methods described below.

*Dataset 1*

15, right-handed, adult subjects (mean age: 35.21 ± 11.48 years, 7 females). Subjects had no history of pain, psychiatric, or neurological disorders. FMRI data were collected while subjects received thermal stimuli across 3 temperatures: 47, 49, and 51 degrees Celsius. Subjects continuously rated, using a finger span device (Apkarian et al., 2001; Baliki et al., 2006), their pain from 0, not painful, up to 100, worst imaginable pain ("pain rating" task.) A control scan was performed while subjects used the finger span device to track a moving bar projected on the screen ("visual rating" task; moving bar replicated for each subject the specific pain rating task temporal pattern). The dataset includes one GLM beta map per subject per stimulus type. The dataset was previously described in (Baliki et al., 2009).

*Dataset 2*

51 healthy, right-handed, adult subjects (mean age: 24 ± 2.29 years, 34 females). Subjects had no history of brain injuries, pain disorders, or psychiatric or neurological diseases. FMRI data was collected while subjects received painful heat stimuli on the right foot dorsum using a $CO_2$ laser, as well as tactile stimuli to the same area using electrical stimulation. Stimuli were not delivered at the same time. Perceived intensities were recorded for every stimulus and only the stimuli with matched perceived intensity for painful heat and touch were selected for GLM analysis. The dataset includes one activation map per subject per stimulus modality – painful heat and touch. The dataset was previously described in (Liang et al., 2019; Su et al., 2019).

*Dataset 3*

14 healthy, right-handed, adult subjects (age: 20 – 36 years old, 6 females). FMRI data were collected while subjects received painful heat stimuli on the right foot dorsum using a $CO_2$ laser, tactile stimuli to the same area using electrical stimulation, visual stimuli using a white disk presented above the right foot, and auditory stimuli delivered via pneumatic earphones. Stimuli were not delivered at the same time. Perceived intensities were recorded for every stimulus and only the stimuli with matched perceived intensity across the four modalities were selected for GLM analysis. The dataset includes one activation map per subject per stimulus modality – painful heat, tactile, auditory, and visual. The dataset was previously described and published in (Liang et al., 2019).

*Dataset 4*

33 healthy, right-handed, adult subjects (mean age: 27.9 ± 9.0 years, 22 females). Subjects had no history of pain, psychiatric, or neurological disorders. FMRI data was collected while subjects received thermal stimuli that varied in one-degree Celsius increments across six temperatures from 44.3 degrees Celsius up to 49.3. Subjects then evaluated each stimulus as warm, and scored it from 0, not perceived up to 99, about to become painful, or as painfully hot, and scored it from 100, no pain, up to 200, worst imaginable pain. The dataset includes an average GLM activation map per subject per stimulus temperature, as well as the corresponding average stimulus ratings. When this dataset was applied dichotomously (pain vs. no pain), we averaged the brain activity maps from the painful and

nonpainful conditions; we omitted subjects who had fewer than two brain activity maps for each condition, resulting in 29 subjects for dichotomous ratings. The dataset was previously described in (Wager et al., 2013; Woo et al., 2015).

*Dataset 5*

14 healthy, right-handed, adult subjects (mean age 22.4 years, range 19–35, 10 females). Subjects had no history of neuropsychiatric disorders, and were not on psychoactive medications. FMRI data was collected while at each trial subjects were presented with a word and had to decide if it refers to a living or nonliving entity. Each word was presented either mirrored or plain. The direction of presented words were interspersed such that we end up with four trial scenarios: Plain-Repeat (PL-RP) where during the trial and the one immediately preceding it, the words were plain; Mirror-Repeat (MR-RP) where during the trial and the one immediately preceding it, the words were mirrored; Plain-Switch (PL-SW) where during the trial the word is plain, and the trial immediately preceding it, the word is mirrored; Plain-Switch (MR-SW) where during the trial the word is mirrored, and the trial immediately preceding it, the word is plain. Data was collected across twelve runs, two training weeks separated two sets of six runs. Dataset includes, up to 12 GLM activation maps (minimum 10) per subject per scenario. The dataset was previously described in (Jimura et al., 2014).

This dataset was provided in subject space. We performed a nonlinear registration of the brains into standard MNI space, 2x2x2 $mm^3$, using FSL FNIRT (Andersson et al., 2007).

*Dataset 6*

213 healthy, adult subjects (mean age 24.1 year (SD = 7.4 year), 101 females). Subjects had no history of physical or mental health condition. fMRI data was collected while subjects performed a voice localizer task. Forty blocks of vocal sounds (20) and non-vocal sounds (20) interspersed with periods of silence were presented while the subjects laid silent and passively listening with their eyes closed in the scanner. This dataset was previously described in (Pernet et al., 2015). Raw fMRI data was downloaded from openneuro.org (https://openneuro.org/datasets/ds000158/versions/1.0.0). fMRI data was preprocessed using FMRIB Expert Analysis Tool (Woolrich et al., 2001), MATLAB and R. Preprocessing included discarding the first 4 volumes for magnetic field stabilization, motion correction, slice-time correction, intensity normalization, band-pass temporal filtering (0.008 Hz – 0.1 Hz), regressing out CSF, white matter, and global signal, censoring of volumes with frame-wise displacement above 0.5 mm, DVARS larger than 2.3, and SD after Z normalization larger than 2.3, as well as two volumes before and two after each censored volume (Huang et al., 2019; Power et al., 2012; Power et al., 2014). We then performed a nonlinear registration of the brains into standard MNI space, 2x2x2 $mm^3$, using FSL FNIRT (Andersson et al., 2007). Task related activation maps (vocal vs silence, and non-vocal vs silence) were derived from a whole brain GLM regression analysis using the FMRIB Software Library (FSL) (Jenkinson et al., 2012; Smith et al., 2004; Woolrich et al., 2009).

### Pain Decoder Templates as candidate across-subject fixed-weight multi-voxel patterns (fwMVPs)

*Neurological Pain Signature (NPS)*

Neurologic Pain Signature, NPS, was shared with us by Tor Wager, whose team developed this across-subject fwMVP (Wager et al., 2013), and has studied its decoding abilities in multiple publications.

*Pain Preferring Voxels (pPV)*

Pain preferring voxels, pPV, is an as-fwMVP decoder developed by Iannetti and colleagues (Liang et al., 2019).

Pain Decoder Template as candidate across-subject fixed-weight meta-GLM multi-voxel pattern

*"Pain" Neurosynth Map (pNsy)*
We used the term-based meta-analysis platform Neurosynth (Yarkoni et al., 2011) to identify a reverse inference brain activity pattern for the term "pain", using association test . We term the obtained pattern as pain-Neurosynth, or pNsy, decoder. Neurosynth uses a probabilistic framework based on Generalized Correspondence Latent Dirichlet Allocation and extracts latent topics from a database of 14,371 published fMRI studies (neurosynth.org, (Yarkoni et al., 2011)). The term "pain" identified 516 studies based on which a brain pattern was generated. The reverse inference association map (FDR corrected <0.01) was used as pNsy, which identifies voxels and their probabilities for being included in the 516 "pain" term associated studies but not in the rest of the >11,000 studies.

*Gaussian Process Decoder*
We used a probabilistic Gaussian Process-based (GP) modeling algorithm (Rasmussen, 2003; Schrouff et al., 2013a) to derive an across-subject fwMVP decoder from the contrast between thermal pain ratings and ratings of visual bars in Dataset 1. We used the publicly available Matlab toolbox PRoNTo (ver2.1.1) (Schrouff et al., 2013a; Schrouff et al., 2013b). We label derived fwMVP decoder pain-GP, or pGP.

Normalized dot product
Throughout this study, we use the normalized dot product (NDP) (eq. 1) as a measure of similarity between templates and brain activation patterns. The NDP is calculated between the vectorized forms of a given decoder template and a stimulus specific brain activation map. The NDP is a scalar between -1 for colinear vectors of opposite direction, and 1 for colinear vectors of same direction. An NDP value of zero means the 2 vectors are orthogonal to each other – no similarity.

$$NDP = T \cdot = \frac{\sum_{i=1}^{n} T_i \cdot \beta_i}{\sqrt{\sum_{i=1}^{n} T_i^2 \cdot \sum_{i=1}^{n} \beta_i^2}} \qquad \text{eq. 1}$$

Where T and β are the vectorized forms of the decoding template and a stimulus specific activation map, $T_i$ and $\beta_i$ are the components of T and β, and n is the number of voxels comprising the brain.

Binary Classification
Two types of binary classifications were performed. The first is a between groups binary classification of brains in painful vs non-painful conditions (or some other decode-comparator pair). We start by calculating the NDP for each brain under each condition; We then use the NDPs as scores to build the Receiver Operator Curve and calculate the area under the curve (AUC). The second classification is a Forced Choice classification, this is a threshold free classification, where the NDP of two brains are compared to each other, and the one with the highest value is classified as "in pain", or as experiencing a higher level of pain than the second brain.

Pattern Smoothing
In order to evaluate the importance of the spatial pattern of fwMVPs on the performance of task classification, NDPs were calculated using spatially smoothed versions of a given decoder. Our hypothesis is that, if a pattern holds task specific information, then spatial smoothing will diminish the

performance of the classifier. Smoothing was done using a 3D isotropic Gaussian kernel filter applied to each template in standard space (eq. 2).

$$T_f(x, y, z|\sigma) = \frac{(T * G(\sigma))(x,y,z)}{(M * G(\sigma))(x,y,z)} \cdot M(x, y, z) \quad \text{eq. 2}$$

Where $T$ and $T_f$ are the original and filtered decoder respectively, $G$ is the Gaussian kernel, $M$ is a binary mask that is True where the decoder is non-zero and False everywhere else, *x, y, z* are voxel coordinates, and $\sigma$ is the kernel standard deviation. The additional $M$ in the numerator resets all non-decoder voxels to zero after filtering – preventing the decoder from bleeding out of its boundary. The convolution in the denominator is the sum of the kernel coefficients where it overlaps with the decoder; this normalization leads to a weighted average using only voxels within the decoder. Together, the additional $M$ in the numerator and the convolution in the denominator correct for boundary effects during filtering. In addition to the original decoder, patterns were progressively smoothed by varying the kernel standard deviation from 1 mm up to 20 mm.

A binarized version of each decoder was also used to simulate a filter with infinite standard deviation, as well as the sign of each filtered decoder at each filter level (sgn($T_f$)), where voxels that are positive become 1, and voxels that are negative become -1, and zero everywhere else. The signed version of the templates was motivated by the fact that in contrast with pPV and pNsy, almost half of the NPS voxels are negative (22,725/47,490), and we needed to investigate the role of sign of the coefficients excluding the effect of the absolute value on decoding. The NDPs generated from these spatial filters were used to calculate the AUC at each smoothing level.

Meta-Analysis

Meta-analysis was performed to obtain average performance estimates for each of the three primary decoders. We modeled each decoder separately since they are 'competing'; as such, the effect of covariance on model parameter estimates is undesirable. Because Dataset 3 contained three comparator tasks, we averaged their performance and estimated the variance of this estimate using the bootstrap technique (1000 replicates); thus, the variance estimate of the average accounts for covariance between the three comparator conditions. No variance stabilizing transformation was performed since the bootstrap distribution of each AUC was approximately normal and transformations provided little gain on average. Both NPS and pNsy were modeled using Datasets 1–4, and pPV was modeled using Datasets 1, 2, and 4, as pPV was derived from Dataset 3. In other words, to use Dataset 4 in the pPV meta-analysis would bias the results in favor of pPV, and we wanted each estimate to be unbiased. We performed a random-effects meta-analysis, fit using restricted maximum likelihood in the *metafor* package using the raw AUCs (Viechtbauer, 2010).

Information Redundancy

We investigated the extent of information redundancy for the three pain the decoders. We wanted to examine whether the spatial extent of a given decoder was needed, and what percent, on average, of the total number of voxels in each decoder was necessary before the classifier performance becomes comparable to the full decoder. Our hypothesis is that if there is no information redundancy, the performance will reach its maximum only when we include the entire decoder; and with increasing redundancy this maximum will be reached with a lower percentage of voxels on average.

Based on the raw, the unfiltered sign, and the infinitely filtered version of each as-fwMVP, we constructed a series of new decoders that included an increasing number of voxels randomly selected from the parent fwMVP without replacement, all remaining voxels were set to zero. We started with ten voxels and increased to the maximum number of voxels in a template. This random sampling was repeated 1,000 time, which produces as many NDPs for each density level. The NDPs were then used to calculate the ROC and its area, which were then averaged to give the average AUC at each percentage level and also calculate associated uncertainty.

<u>Voxel Weights</u>
We investigated whether or not voxels with higher coefficients (in absolute value) encode more state specific information compared to voxels with lower coefficients. To address this question, we binned each fwMVP voxels by their absolute weights, such that the top 10% of absolute voxel weights were in the first bin, the second 10% were in the second bin, etc., and built a decoder from each tier. We then used those templates to calculate the NDPs and the AUCs as a function of voxel coefficient tier. In addition to the 10% bin width and unfiltered decoders, we also generated decoders using bin widths of 1%, 5%, and 20%, as well as decoders from the sign of the unfiltered, and infinitely filtered versions.

<u>Role of Brain Areas</u>
We investigated whether decoder voxels from certain brain regions perform better than others. We selected pNsy as the decoder for this analysis given the probabilistic meaning of its voxel weights. We thresholded the decoder (voxel weights z > 6) and generated a new decoder from each distinct cluster; we ended up with seven new decoders. We then evaluated the pain decoding performance of each new decoder on datasets 1 to 4. we applied a gaussian spatial filter (sd = 10 mm) before thresholding, otherwise we end up with too many fragmented clusters.

<u>Bayesian classification for identification</u>
We created a nonparametric Bayesian classification model to probabilistically classify subjects as being in a certain state given their brain activity map. This model was trained and run on all subjects across all pain Datasets (Fix and Hodges, 1951; Silverman, 1986), in addition to the voice dataset (Pernet et al., 2015).

Starting with the pain datasets, we started with a matrix containing all subjects, tasks, and their respective normalized dot products (NDP). Each subject was sampled one at a time. Using the remaining subjects, a probability density functions (pdf) of normalized dot products was created for each task. To create these pdfs, we used kernel density estimation with a Gaussian kernel and a bandwidth chosen using the Sheather-Jones method (Sheather and Jones, 1991). Specifically, a pdf was created for each of the comparator conditions (visuomotor, touch, audition, vision, and nonpainful heat) and pain. All of the pain conditions were modeled as one distribution, as a tacit assumption of these decoders is that "physical pain" is a single construct. From these distributions, we could calculate a posterior probability, P(pain | NDP), for each individual *i*:

$$P(pain|NDP_i) = \frac{\hat{f}_{pain}(NDP_i|pain)P(pain)}{\sum_{j=1}^{k}\hat{f}_j(NDP_i|task_j)P(task_j)} \qquad \text{eq. 3}$$

where $\hat{f}_{pain}(NDP_i)$ and $\hat{f}_j(NDP_i)$ are the kernel density estimates used for $NDP_i$ (i.e., derived from all other brains) in pain or task $j$ (where tasks $j$ = 1, …, $k$ include all comparator tasks and pain). Priors, $P(\cdot)$, were derived from the number of studies in Neurosynth that contains:

- "pain" = 516
- "tactile" OR "touch" = 110 + 225
- "visually" OR "vision" = 333 + 137

- "auditory" = 1253
- "visuomotor" = 153
- "heat" (from old Neurosynth) = 61

All study counts were obtained on December 10, 2019. Because they were obtained from Neurosynth and each study is given equal weight, the priors assume an equal number of subjects across studies, and thus estimates the probability of a brain undergoing each of these tasks in the "neuroimaging world," if we consider these tasks to be the neuroimaging world. Of note, these priors provided more optimistic estimates as compared to uniform priors.

For both NPS and pNsy, all subjects were used to obtain the posterior distribution. However, to obtain an unbiased posterior distribution for pPV, we did not include subjects from Dataset 3 (i.e., from which pPV was derived).

This process was repeated for the voice test dataset (106 subjects). However, because the tasks in the voice dataset were unique, we used a flat prior (i.e., prior probability = ½ for each of the two tasks).

Calculation of distributional overlap for identification

We calculated the overlap between the distributions of decodee and comparator NDPs as a marker of identifiability. The overlapping region of probability density functions contains information that cannot be used to identify; thus, lower overlap corresponds to higher identifiability. To calculate overlap, we first fit each NDP distribution (e.g., NPS pain and NPS nonpain, separately) using kernel density estimation with an Gaussian kernel and a bandwidth chosen using the Sheather-Jones method (Sheather and Jones, 1991). We then had $\hat{f}_{decodee}(\cdot)$ and $\hat{f}_{comparator}(\cdot)$, kernel density estimates for the decodee and comparator, respectively. We integrated over their minimum to calculate their overlap:

$$\int_{-1}^{1} \min\left(\hat{f}_{decodee}(x), \hat{f}_{comparator}(x)\right) dx$$

Normalized Dot Product – Stimulus Relationship

In this analysis we wanted to investigate the relationship between the NDP and stimulus rating as well as stimulus intensity. Dataset 4 includes information about stimulus intensity and stimulus rating. We fit the data using locally estimated scatterplot smoothing (LOESS) (Cleveland and Devlin, 1988).

Within Study vs. Across Study Decoders

Given that pNsy is based on a meta-analysis of study-level GLM brain activity maps, we created decoders from four datasets by averaging subject-level GLM brain activity maps obtained from a pain task. These study-level decoders were then used to classify brains as pain vs. no pain, in accordance with the task.

```
for each study i
        average beta maps for the pain task in study i
end

for each study i
        for each study j
                for each subject k in study j
```

*for each task l in subject k*
*calculate cosine similarity of between TASK_lk and DECODER_i*
*end*
*end*

*calculate AUC for DECODER_i applied to STUDY_j*
*end*
*end*

Within Subject vs Across Subject Decoders

Given the variability of fMRI data, both within-subject and across-subject, we wanted to answer the following question: will decoding mental states of a particular subject using a template derived from data of the same subject be more accurate than decoding of mental states of a group of subjects using a template derived from the group's data? Are within-subject decoders superior to between-subject decoders? The following analysis addresses this question using Dataset 6.

*Within Subject decoding*

Below is a pseudo-code for the within subject analysis.

*for each subject i*
*for each task j*
*randomly select half the task j beta maps replicates,*
*average voxel-wise to get inter subject i, task j specific decoding template Tj,*
*label the remaining task j replicates as TASK_j,*
*calculate SRs of each beta map in TASK_j using Tj,*
*for each task k ≠ j*
*randomly select half the task k beta maps replicates and label as TASK_k,*
*calculate SRs of each beta map in TASK_k using Tj,*
*calculate AUC for correctly classifying TASK_j and TASK_k beta maps,*
*end*
*end*
*end*
*Average the AUCs along all subjects,*
*repeat from the start 1,000 times.*

This will result in average AUC estimates for the classification of each possible task pairs (i,j) using both $T_i$ and $T_j$. All performed within-subject.

*Between Subject decoding*

Below is a pseudo-code for the between subject analysis.

*for each subject i*
*for each task j*
*average beta map replicates to get one beta map per subject per task to form the between-subjects dataset,*
*end*
*end*

*for each task j*

*randomly select half the task j beta maps (from the between-subjects dataset),*
*average voxel-wise to get between-subjects task j specific decoding template Tj,*
*label the remaining task j replicates as TASK_j,*
*calculate SRs of each beta map in TASK_j using Tj,*
*for each task k ≠ j*

*randomly select half the task k beta maps replicates and label as TASK_k,*
*calculate SRs of each beta map in TASK_k using Tj,*
*calculate AUC for correctly classifying TASK_j and TASK_k beta maps,*

*end*

*end*

*repeat from the start 1,000 times.*

This will result in AUC estimates for the classification of each possible task pairs (i,j) using both $T_i$ and $T_j$. All performed between-subject.

Decoders derived from Dataset 5 and Dataset 6

We created fwMVP decoders from Dataset 5 (Jimura et al., 2014) and Dataset 6 (Pernet et al., 2015) to assess the generalizability of our results to other cognitive domains. Four approaches were used to create these decoders: Support Vector Machine, LASSO-PCR, Gaussian Process Classification, as well a GLM contrast of activation maps. Training and testing of the decoders were similar across all four approaches, with some minor differences in the treatment of each dataset in how we select the training and testing groups. Assuming we have our training and testing groups, the procedure is as follows:

1. *Perform a second-level group analysis with cluster-based thresholding corrected for multiple comparisons by using the null distribution of the maximum cluster mass (FSL randomize (Woolrich et al., 2004), option -C) on the training group for the contrast GLM activation maps mr_rpt > (mr_sw, pl_rp, pl_sw) for Dataset 6, and vocal_sound > non-vocal for Dataset 7.*
2. *Binarize the group contrast map; this will be the mask of voxels of interest for building our decoders.*
3. *Use SVM, LASSO-PCR, Gaussian Process to generate the decoder with the activation maps (GLM) of the training group. For GLM decoders, the mean difference in activation maps within this same masked region was used.*
4. *Perform the normalized dot product of the decoder with the activation maps in the testing group to calculate the signature response and calculate the AUC of the classification exercise.*

Dataset 5 include several replicate activation maps per task for each of the 14 subjects, we preprocessed the data as follows:

1. *Average all task replicates for each subject.*
2. *Randomly split the subjects into two seven subject groups: training and testing.*
3. *Create a template and test it as described above.*
4. *Repeat 100 times from step 2 and build the AUC distribution.*

After preprocessing, Dataset 7 included 213 subjects and had one activation map per stimulus per subject. The large number of subjects allows us to split it into a training group (107 Subjects), and a validation group (106 Subjects) without the need for permutation. Because the sample was large, we calculated the AUC confidence interval using normal assumptions of a binomial proportion (eq. 4)

$$95\%\ CI = p \pm 1.96\sqrt{\frac{p(1-p)}{n}} \quad \text{eq. 4}$$

where $p$ is the AUC.

*SVM and Gaussian Process*

We used the Matlab toolbox PRoNTo (ver2.1.1) (Schrouff et al., 2013a; Schrouff et al., 2013b) to derive the decoders using SVM (Cristianini and Shawe-Taylor, 2000; Mourao-Miranda et al., 2007), and Gaussian Process (Rasmussen, 2003; Schrouff et al., 2013b). Data was split into two groups; Group 1 included activation maps of the mr_rp task for Dataset 5, and of the vocal_sound stimulus for Dataset 6; Group 2 included the activation maps of the mr_sw, pl_rp, and pl_sw for Dataset 5, and non-vocal_sound for Dataset 6. All maps were input as independent datapoints. We performed a binary classification analysis and used "Binary Support Vector Machine" for SVM, and "Binary Gaussian Process Classifier" for gaussian process, and constrained the analysis to voxels within the mask created from the second-level group analysis

*LASSO-PCR* was used to generate decoders following the methods described by Wager *et al.* (Wager et al., 2011; Wager et al., 2013) and was implemented in R. An $n \times p$ sparse matrix of subjects ($n$) and voxels ($p$) was column-wise centered and scaled. Of note, sparse columns were left sparse since their scaled estimates are undetermined. Principal components analysis was performed using singular value decomposition on the column-scaled matrix to obtain a new $n \times n$ predictor matrix, $\mathbf{X}_{\mathbf{PCA}}$, and a $p \times n$ rotation matrix, **R**. The reduced predictor matrix, $\mathbf{X}_{\mathbf{PCA}}$, was used in a logistic regression with $L_1$ regularization (LASSO) (Efron et al., 2004; Friedman et al., 2010; Simon et al., 2011). Hyperparameter $\lambda$ was chosen to minimize binomial deviance using leave-one-out cross-validation across 1,000 $\lambda$'s; default glmnet parameters were used to determine the exact grid range. An $n \times 1$ vector of penalized coefficients was pre-multiplied by rotation matrix **R** to obtain a $p \times 1$ vector of voxel weights. This vector of voxel weights served as the decoder.

*GLM* was used to generate contrast-based decoders. These simply used the average difference between unsmoothed GLM activity maps (*e.g.*, mean(vocal) – mean(non-vocal)), masked to the same thresholded region as the other decoders.
