## Supplemental Results for "The Hard Limits of Decoding Mental States: The Decodability of fMRI"

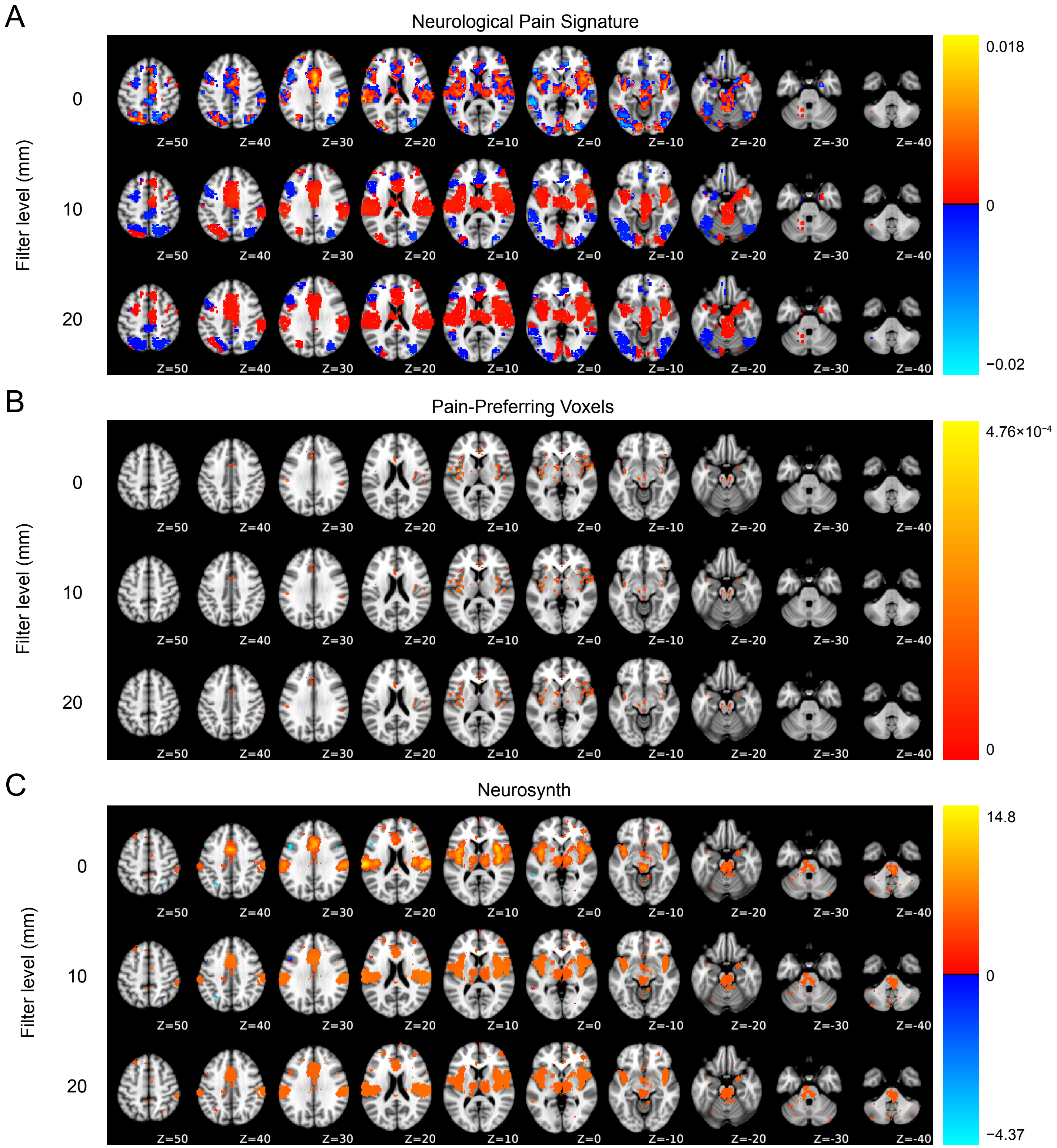


**Supplemental Figure 1. Spatial extents and influence of Gaussian spatial filtering for the three pain decoders. A.** The fwMVP Neurological Pain Signature (NPS), **B.** The fwMVP Pain Preferring Voxel (pPV), **C.** The meta-GLM Pain-Neurosynth (pNsy). Presented are dorsal slices from Z=50 to Z=-40 in standard MNI space. Top row in each panel shows the raw, unfiltered decoder. Row 2, shows resultant decoder pattern after 3D spatial smoothing using a Gaussian kernel with a standard deviation of 10 mm (5x voxel width); Row 3, decoder pattern after Gaussian smoothing with a standard deviation of 20 mm (10x voxel width). Note, the differences in spatial patterns and extent between decoders: NPS covers a larger extent of the brain than pPV or pNsy; NPS voxels outside of pNsy are mostly composed of negative values; and pPV is comprised of sparse voxels within the bounds of pNsy. Moreover, the positive weights of all three decoders sample mostly common brain regions (bounded by pNsy). For all three decoders, spatial variability in voxel weights decreases with increasing size spatial filtering: brain regions become more homogeneous; regions with both positive and negative coefficients become either positive or negative with filtering; eventually, at infinity, all template coefficients become a single positive value (at infinite filtering decoders only contain location information: binary decoders: 0, 1). Note, due to the sparsity of pPV, filtering effects are subtle.

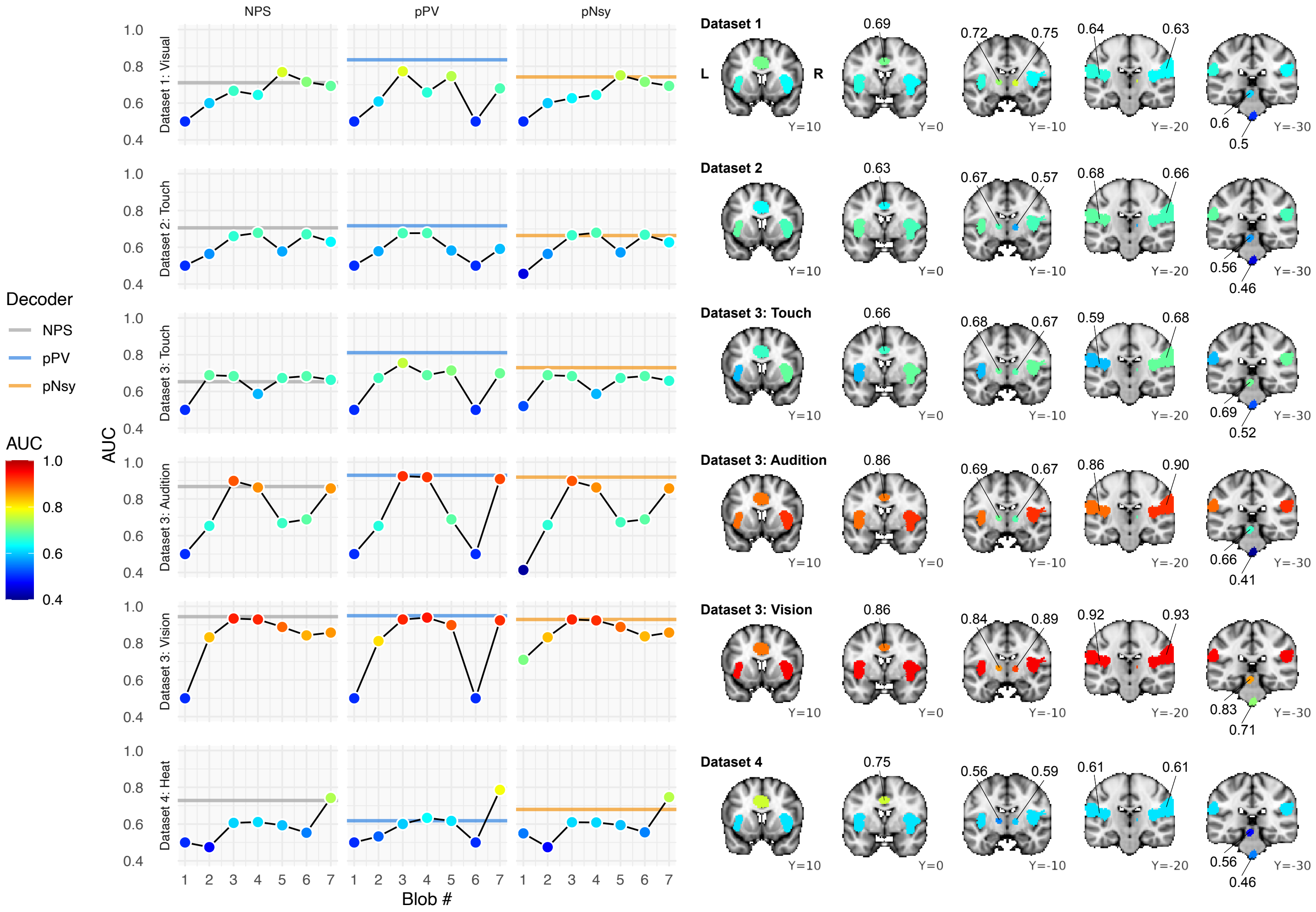


**Supplementary figure 2. Discrimination performance for different clusters to assess dependence on specific brain regions.** There is a long debate in the literature regarding which brain regions show better specificity to nociceptive stimuli and/or pain states. Here we address the issue from a decoding viewpoint. Here we ask the question whether there are privileged locations for decoding pain. As all three pain decoders (NPS, pPV, and pNsy) overlap most with pNsy, we used the latter to create separate blobs, within each of which we tested the performance of all three decoders. Pain-Neurosynth was thresholded at $z = 6$ to obtain 7 distinct and contiguous regions, or "blobs" (locations illustrated on standard atlas on the right)." NPS, pPV, and pNsy values in each of the 7 blobs were used as separate decoders, and their respective performance was obtained for discriminating pain from comparator states across all studies. Three left columns show discrimination performance for each decoder, within each blob, for all tasks as indicated. The right brain maps show performance for pNsy, localized to each blob (colors match the color distribution in the right pNsy column), for all four task comparisons.

**Dataset 1: Visual**, painful heat with continuous rating vs visual rating (15 subjects). **Dataset 2: Touch**, painful heat vs tactile stimuli (51 subjects). **Dataset 3: Touch**, Painful heat vs tactile stimuli (14 subjects). **Dataset 3: Auditory**, Painful heat vs auditory stimuli (14 subjects). **Dataset 3: Visual**, Painful heat vs visual stimuli (14 subjects). **Dataset 4: Heat**, Painful heat vs non-painful warm thermal stimuli (33 subjects).

Our performance metric is the area under the receiver operating characteristic curve (AUC).

For any given study, multiple blobs for multiple decoders performed equally well and matched the performance of the full decoder (grey, blue, and orange lines). This result suggests lack of any given blob showing consistently better specificity than others. Some blobs in isolation performed better than the entire decoder (grey, blue, and orange lines), but the effect was study- and decoder- specific. In some instances (Blob #1 with NPS and pPV, Blob #6 with pPV), the Blobs had no spatial overlap with the decoders; for these cases, the blob received an AUC of 0.5. Blob 1 (inferior brainstem) was the region most consistently showing worst decoding ability across studies and decoder types. This is partially explained by the exclusion of Blob #1 in NPS and pPV; but in pNsy, we suspect this effect is due to the influence of physiological noise that contaminates brainstem activity.

Blob 1 = inferior brainstem; blob 2 = superior brainstem; blob 3 = right insula & parietal lobe; blob 4 = left insula & parietal lobe; blob 5 = left thalamus; blob 6 = right thalamus; and blob 7 = anterior cingulate cortex.

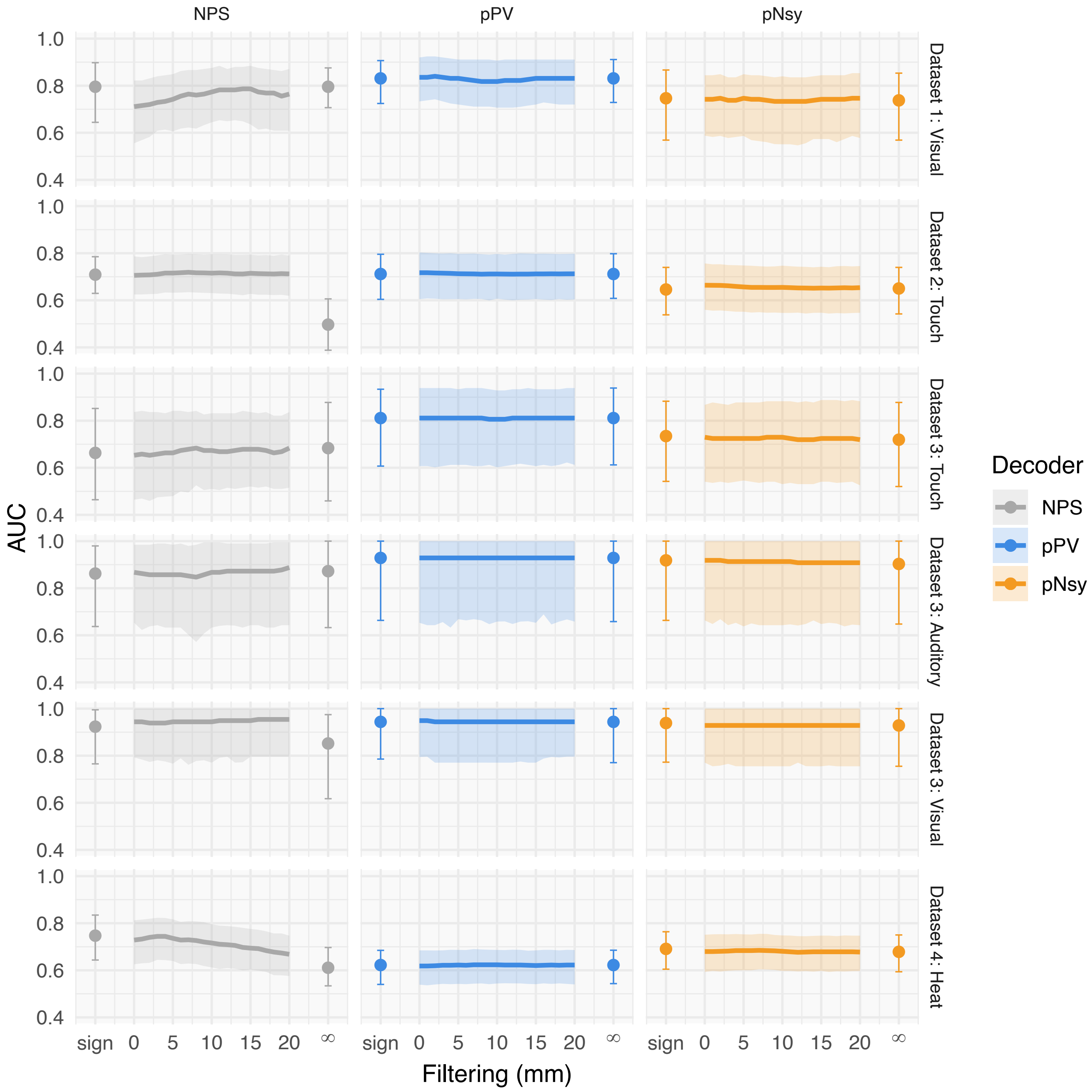


**Supplemental Figure 3. Discrimination performance as a function of manipulating weights of decoders.** We examined relative to raw decoders, preserving sign only (-1, 0, +1) (sign), increasing size spatial smoothing (0-20 mm), and infinite spatial smoothing (0, +1) (∞), for the *original unfiltered* three pain decoders (NPS, pPV, and pNsy), examined in four studies contrasting pain states with various control conditions. **Dataset 1: Visual**, painful heat with continuous rating vs visual rating (15 subjects). **Dataset 2: Touch**, painful heat vs tactile stimuli (51 subjects). **Dataset 3: Touch**, Painful heat vs tactile stimuli (14 subjects). **Dataset 3: Auditory**, Painful heat vs auditory stimuli (14 subjects). **Dataset 3: Visual**, Painful heat vs visual stimuli (14 subjects). **Dataset 4: Heat**, Painful heat vs non-painful warm thermal stimuli (33 subjects).
Our performance metric is the area under the receiver operating characteristic curve (AUC). Values at "sign" are AUC when only the sign of the decoder is preserved (-1, 0, +1); 0 filtering are AUC using raw unfiltered decoders; curves are plots of AUC vs kernel standard deviation for 0-20 mm filtering; ∞ in each graph are performances using filtered templates with infinite standard deviation (binarized templates; 0, +1); whiskers and shaded area are 95% confidence intervals estimated using the bias-corrected and accelerated bootstrap.
Notice that, except for NPS, decoding performance is essentially unaffected by spatial filtering even with an infinitely wide filter which completely deletes the pattern. Generally, these findings suggest that the information captured by all three pain decoders is not dependent on the fine-grained patterns of weights that constitute the decoders.
Performance significantly degrades for infinity-filtered NPS in Dataset 2: Touch (**Row 2**), and slightly for Dataset 4: Heat (**Row 6**). Even in these cases the NPS sign-only shows the same performance as raw NPS, revealing that the degradation is due to assigning positive weights to brain regions where activity is not related to the pain task.

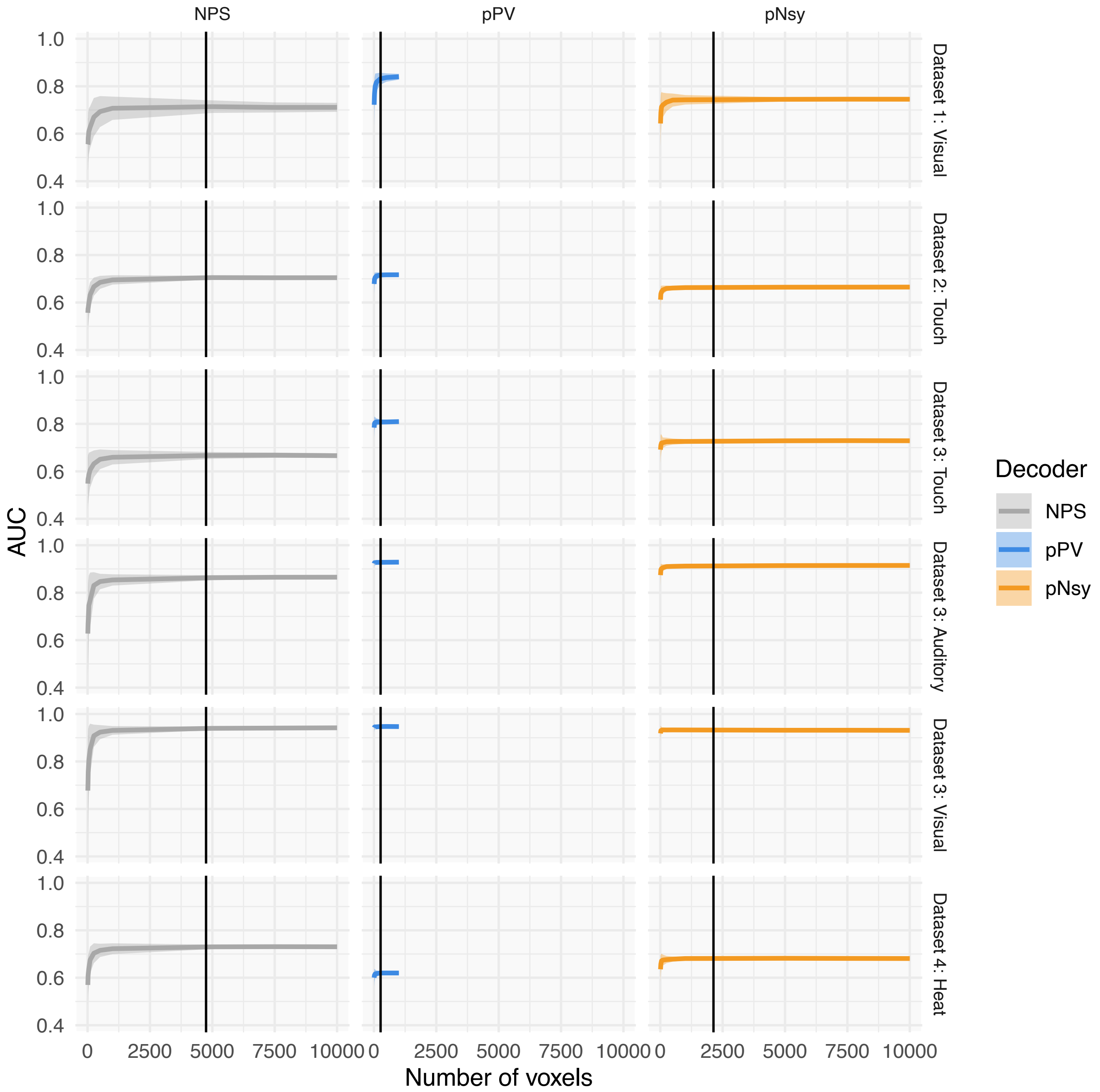


**Supplemental Figure 4. Testing for information redundancy by random subsampling of *unfiltered* decoders.** For all three *unfiltered* decoders (NPS, pPV, and pNsy; each column), an increasing number of voxels was selected at random to form new decoders. The new decoders were comprised of voxels from 10 up to the total number of voxels in the parent template. Discrimination performance was measured using each new constructed decoder.

**Dataset 1: Visual**, painful heat with continuous rating vs visual rating (15 subjects). **Dataset 2: Touch**, painful heat vs tactile stimuli (51 subjects). **Dataset 3: Touch**, Painful heat vs tactile stimuli (14 subjects). **Dataset 3: Auditory**, Painful heat vs auditory stimuli (14 subjects). **Dataset 3: Visual**, Painful heat vs visual stimuli (14 subjects). **Dataset 4: Heat**, Painful heat vs non-painful warm thermal stimuli (33 subjects).

Performance metric is the area under the receiver operating characteristic curve (AUC). Shaded areas are standard deviations due to resampling voxels and thus are insensitive to the number of permutations.

We note that the decoding performance reaches that of its full version very quickly. Less than 10% of the parent decoder was required to reach peak performance (black bar marks).

These results indicate existence of sufficient redundancy that on average only a random 10% of the patterns of voxels comprising any of the *unfiltered* decoders was needed to achieve best performance, for all task discriminations.

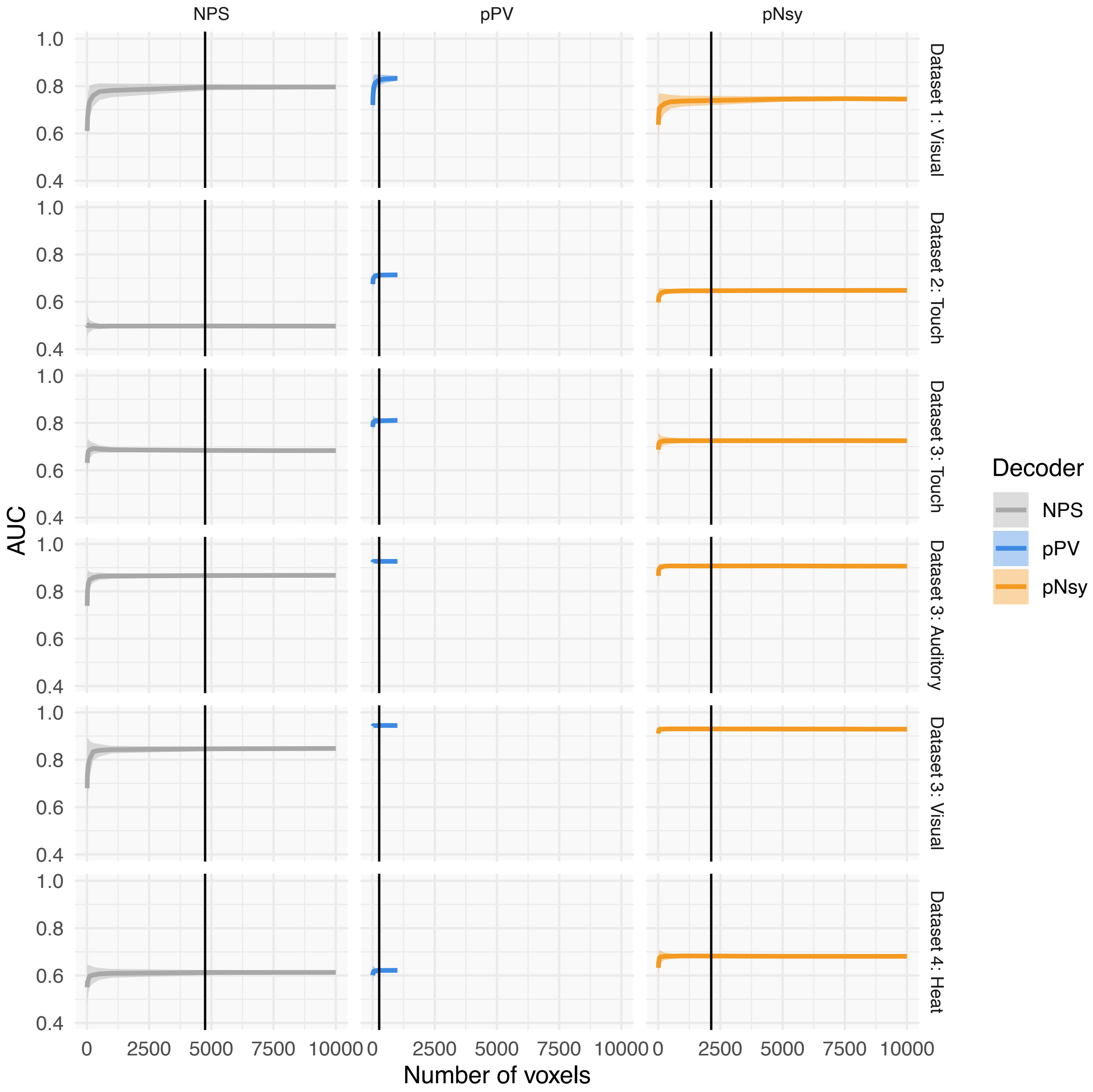


**Supplemental Figure 5. Testing for information redundancy by random subsampling of *infinitely-filtered* decoders.** For all three *infinitely filtered* (0,1) decoders (NPS, pPV, and pNsy; each column), an increasing number of voxels was selected at random to form new decoders (from 10 voxels to the full template). Classification performance was measured using each new decoder template. Shaded areas are standard deviations due to resampling voxels and thus are insensitive to the number of permutations.
**Dataset 1: Visual**, painful heat with continuous rating vs visual rating (15 subjects). **Dataset 2: Touch**, painful heat vs tactile stimuli (51 subjects). **Dataset 3: Touch**, Painful heat vs tactile stimuli (14 subjects). **Dataset 3: Auditory**, Painful heat vs auditory stimuli (14 subjects). **Dataset 3: Visual**, Painful heat vs visual stimuli (14 subjects). **Dataset 4: Heat**, Painful heat vs non-painful warm thermal stimuli (33 subjects).
Similar to using unfiltered templates, we note that the decoding performance reaches peak performance with 10% of the parent template (black bars). These results show that only location information of a random 10% of the voxels comprising any of the decoders was sufficient to achieve best performance, in all task discriminations.

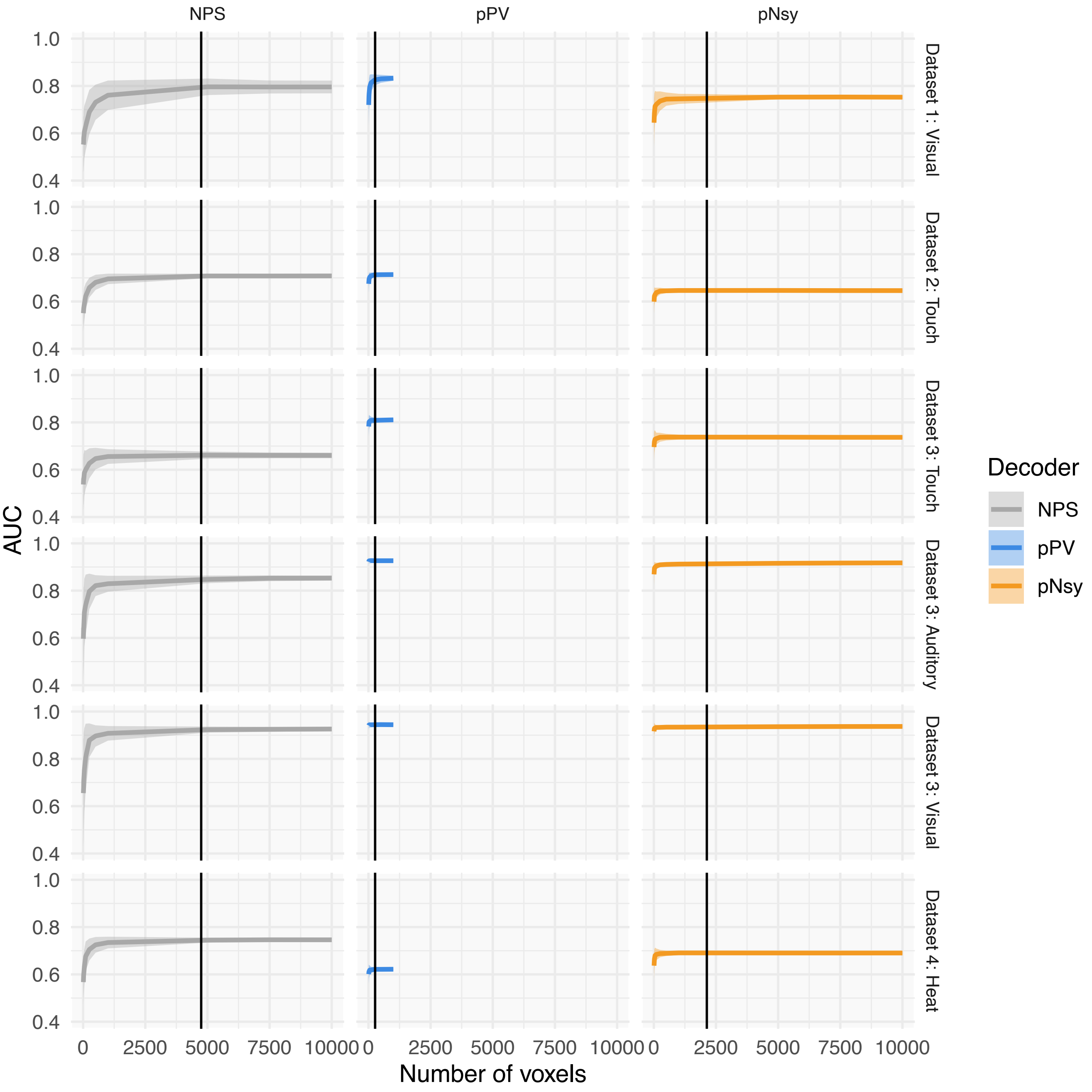


**Supplemental Figure 6. Testing for information redundancy by random subsampling of *sign preserving* decoders.** For each *sign-preserving* decoder (original templates reduced to three values: -1, 0, +1), an increasing number of voxels was selected at random to form new decoders (from 10 voxels to the full template). Discrimination performance was measured using each new decoder template. Shaded areas are standard deviations due to resampling voxels and thus are insensitive to the number of permutations. **Dataset 1: Visual**, painful heat with continuous rating vs visual rating (15 subjects). **Dataset 2: Touch**, painful heat vs tactile stimuli (51 subjects). **Dataset 3: Touch**, Painful heat vs tactile stimuli (14 subjects). **Dataset 3: Auditory**, Painful heat vs auditory stimuli (14 subjects). **Dataset 3: Visual**, Painful heat vs visual stimuli (14 subjects). **Dataset 4: Heat**, Painful heat vs non-painful warm thermal stimuli (33 subjects).

We again note that the decoding performance reaches peak performance with 10% of the *sign-preserving* decoders (black bars).

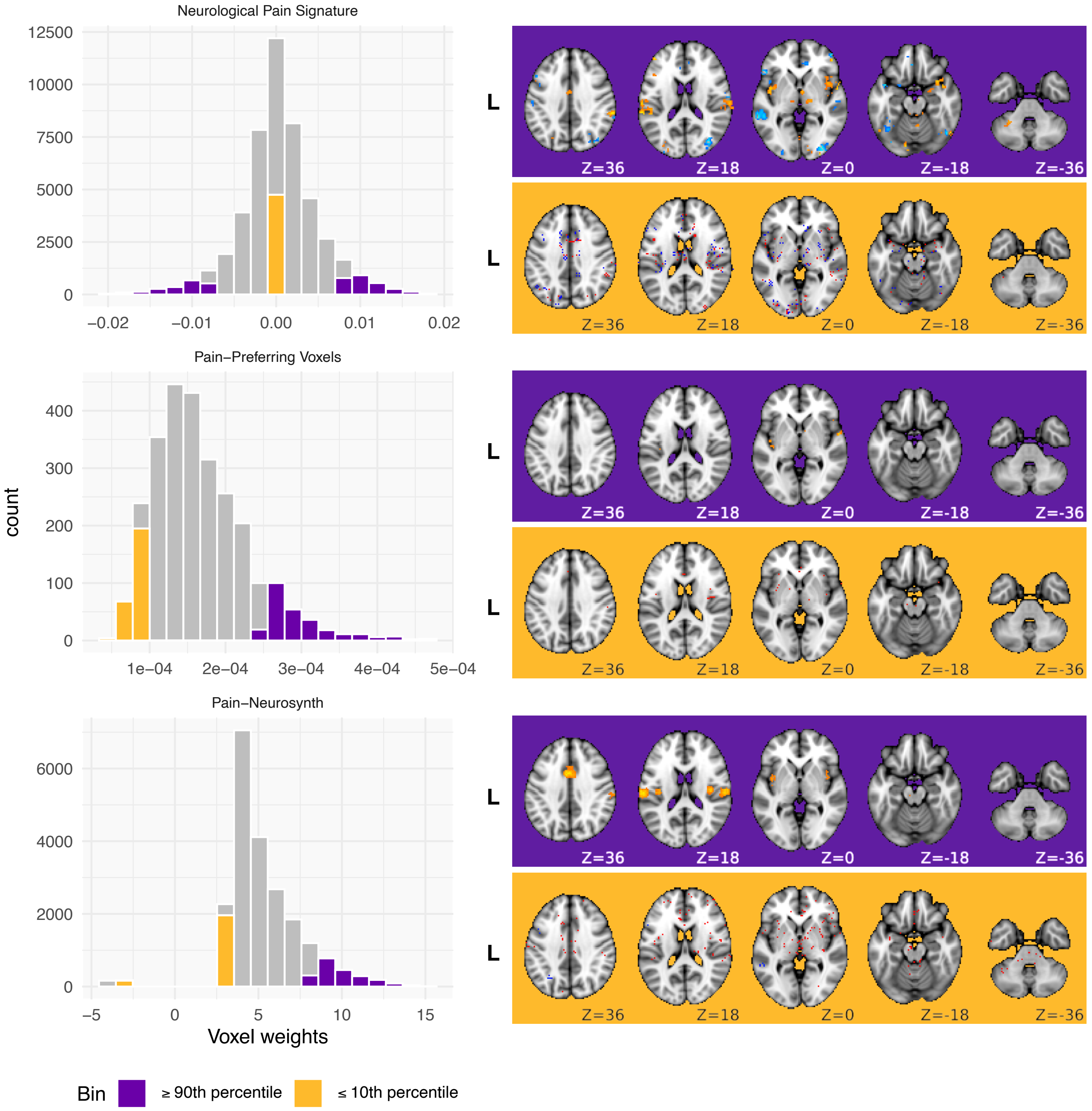


**Supplementary Figure 7. Weight-ordered thinning of decoders.** The figure illustrates procedure used for weight-ordered thinning. For a given distribution of weights, new decoders were constructed from the absolute value of their coefficients using different bin widths (1%, 5%, 10%, and 20% of the total number of voxels in each decoder). The approach systematically quantifies the contribution of different coefficient-weighted brain regions to discrimination performance. All three pain decoders at least imply the importance of different brain regions based on associated coefficients. In NPS and pPV, brain regions with highest positive and lowest negative coefficients should classify pain better than locations with lower absolute coefficients. In pNsy, weights reflect confidence of discriminating "pain" term associated studies from all other studies included in Neurosynth (association test). Therefore, we used weight ordered thinning to examine differential contribution of the weights to discrimination performance. Results are shown in **Supplementary figures 7-9**.

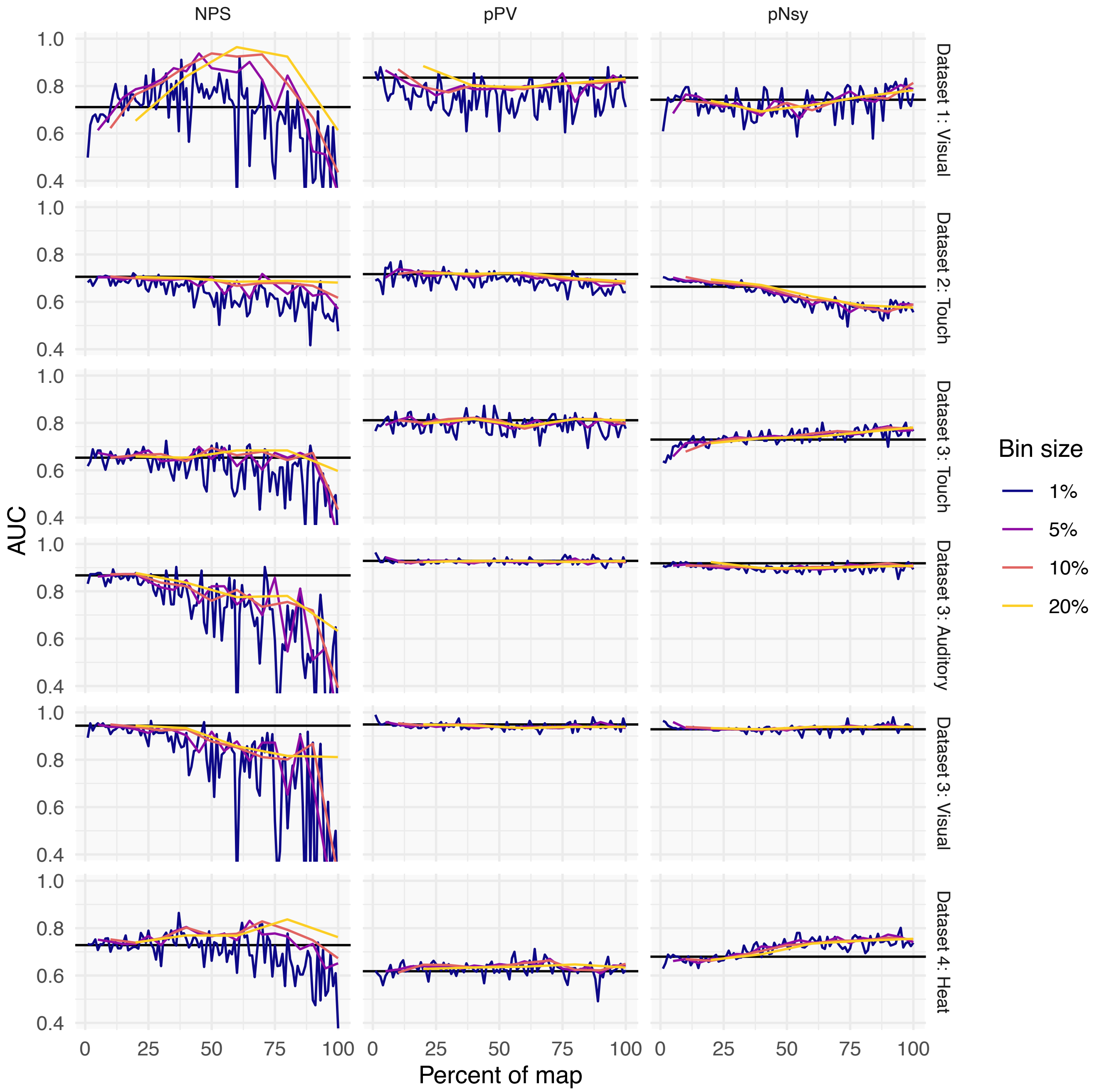


**Supplemental Figure 8. Weight-ordered thinning of *unfiltered* decoders.** For each decoder (NPS, pPV, pNsy), we binned the voxels according to the absolute value of their coefficients. We used four bin widths: 1%, 5%, 10%, and 20% of the total voxel numbers. We then built a decoding template from each binning, and for eac*h* bin width (**Supplementary figure 7**). X-axis of the plot represent the bin number (percent map), Y-axis is performance (AUC), full decoder performance plotted as black line.
**Dataset 1: Visual**, painful heat with continuous rating vs visual rating (15 subjects). **Dataset 2: Touch**, painful heat vs tactile stimuli (51 subjects). **Dataset 3: Touch**, Painful heat vs tactile stimuli (14 subjects). **Dataset 3: Auditory**, Painful heat vs auditory stimuli (14 subjects). **Dataset 3: Visual**, Painful heat vs visual stimuli (14 subjects). **Dataset 4: Heat**, Painful heat vs non-painful warm thermal stimuli (33 subjects).
If weight distribution was an important determinant for performance then we would expect an inverse relationship between AUC and bin number. Instead we observe mostly an invariant relationship, especially for pPV and pNsy decoders. These findings agree and complement the information redundancy results; demonstrating that the values of voxel coefficients have a minimal and inconsistent impact on decoder performance.
However, the unfiltered NPS shows a weak inverse relationship between AUC and bin number for all datasets and comparator stimuli, most apparently at the smallest bin width (1%) and for bins with absolute values in the lowest 50%. Note when we perform the same analysis now using signed-unfiltered decoders, with binning based on their original weights, the results essentially replicate this figure (data not shown). The latter again refutes, even for NPS, the importance of weight distribution on performance, and instead highlights the fact that in NPS location of negative weights has a small contribution on discrimination performance.

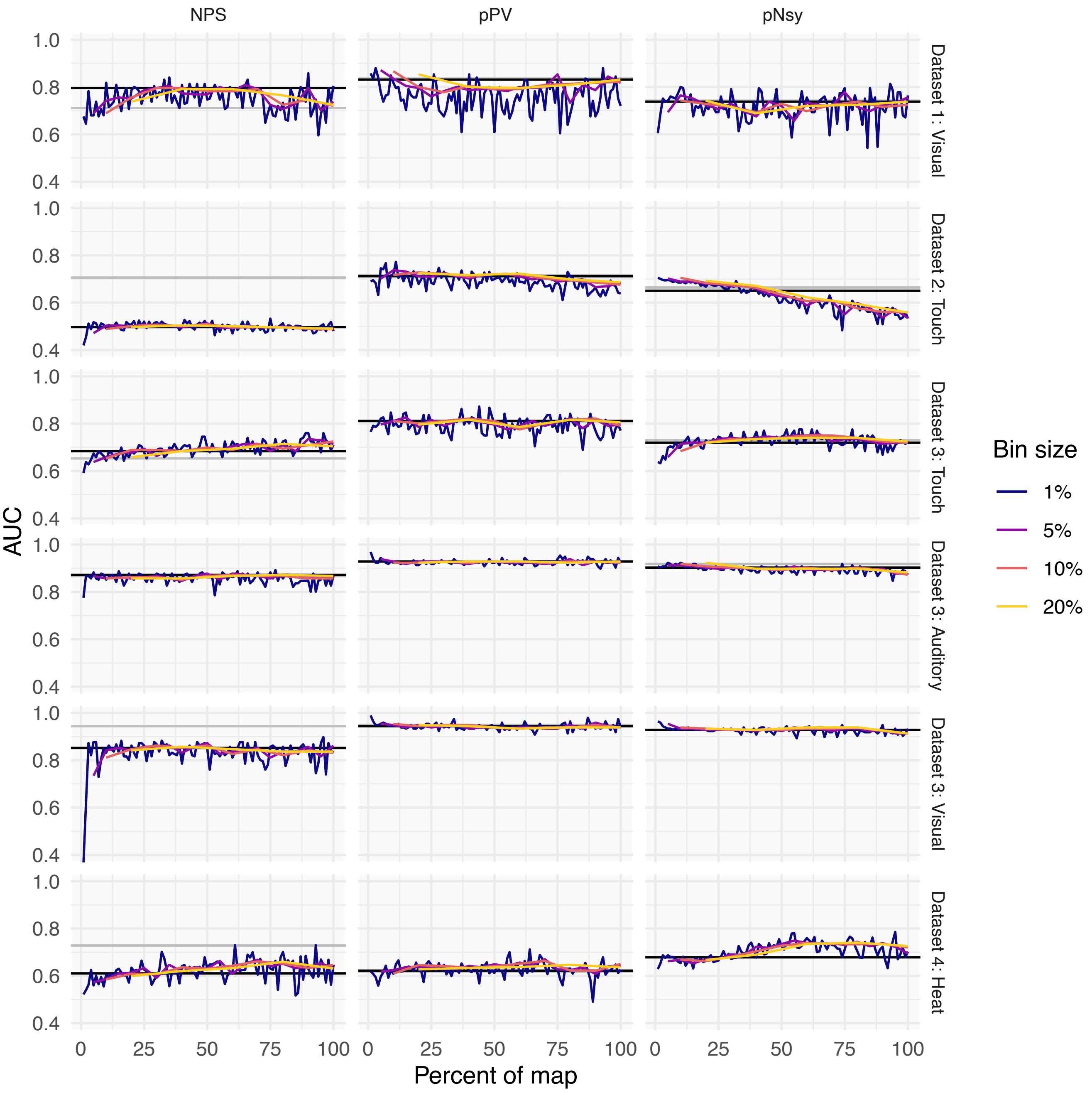


**Supplemental Figure 9. Weight-ordered thinning of *infinitely-filtered* decoders.** For each *infinitely-filtered* (0, +1) decoder (NPS, pPV, pNsy), we binned the voxels according to the absolute value of their original unfiltered coefficients. We used four bin widths: 1%, 5%, 10%, and 20% of the total voxel numbers. We then built a decoding template from the locations for each bin, and for each bin width (**Supplementary figure 7**), but now only using infinitely-filtered decoders (location only: 0, +1). X-axis of the plot represent the bin number (percent map), Y-axis is performance (AUC), full decoder performance plotted as black line.
**Dataset 1: Visual**, painful heat with continuous rating vs visual rating (15 subjects). **Dataset 2: Touch**, painful heat vs tactile stimuli (51 subjects). **Dataset 3: Touch**, Painful heat vs tactile stimuli (14 subjects). **Dataset 3: Auditory**, Painful heat vs auditory stimuli (14 subjects). **Dataset 3: Visual**, Painful heat vs visual stimuli (14 subjects). **Dataset 4: Heat**, Painful heat vs non-painful warm thermal stimuli (33 subjects).
Strikingly we observe no effect of binning on performance, and at all binnings performance matches that seen for the unfiltered original decoders. Therefore, locations of distinct weightings have no tangible preferential added value to overall performance of these decoders.

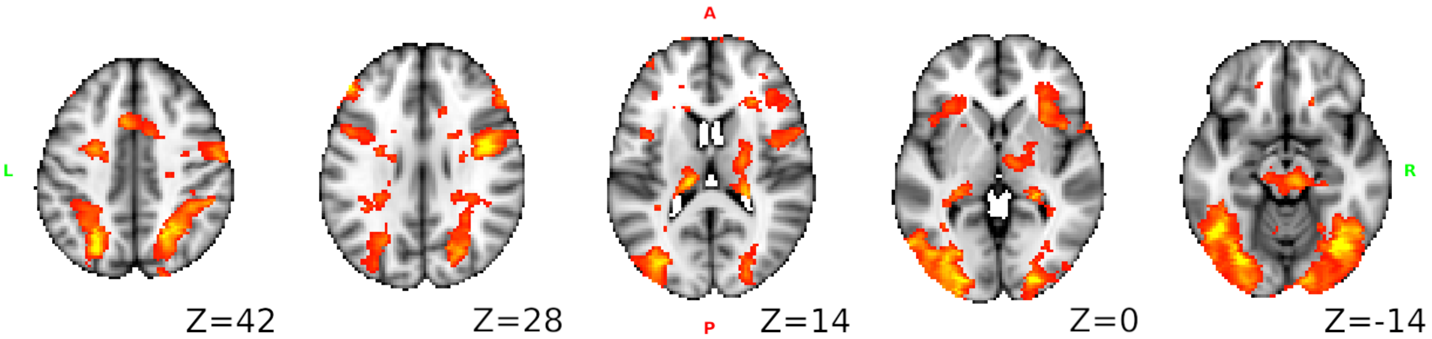


**Supplemental Figure 10. GLM contrast map for Dataset 5** (Jimura et al.) **(*1*).** For this dataset, we had access to individual subjects' GLM analyzed activity maps in native space. Thus, the data had to be registered into standard space prior to using it in discrimination and identification analyses, and prior to generating various types of decoders. Therefore, it was imperative to ascertain that the manipulations we had imposed on the dataset still resulted in activity consistent with published results. Using the registered data from Jimura et al., we replicated their results for "mirror-repeat" versus "plain-repeat" task contrast using *randomise* in FSL (z-values, thresholded above 2.3). Our resulting contrast map closely resembles that of Jimura et al. (*1*).

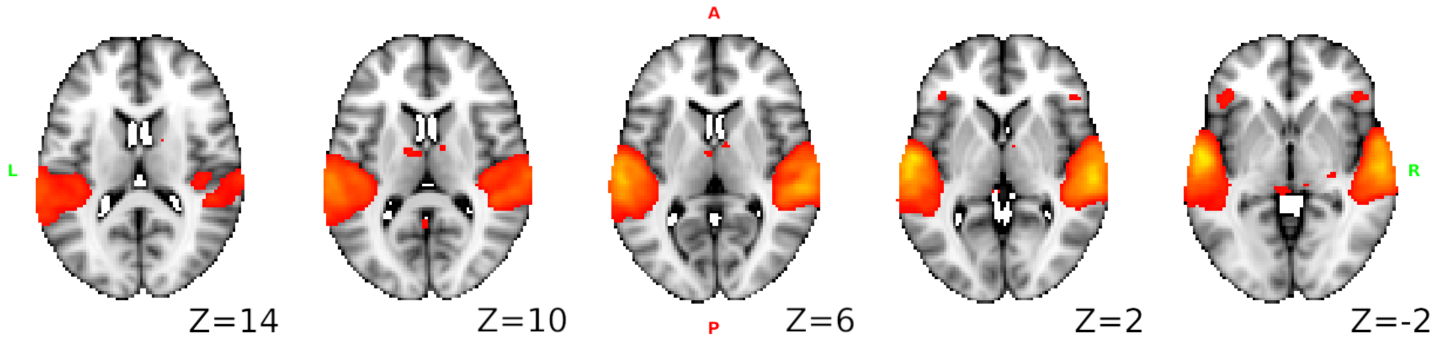


**Supplemental Figure 11. GLM contrast maps for Dataset 6 (Pernet et al.) (*2*).** This dataset was only available is raw time series. We, thus, performed standard preprocessing and post-processing to generate individual subject activity maps in standard space. Therefore, it was imperative to ascertain that the manipulations we had imposed on the dataset still resulted in activity consistent with published results. Using the raw data and after our pre- and post-processing, we replicated their results for "vocal" versus "non-vocal" task contrast using *randomise* in FSL (z-values above 2.3). Our resulting GLM contrast map closely resembles that of Pernet et al. (*2*)

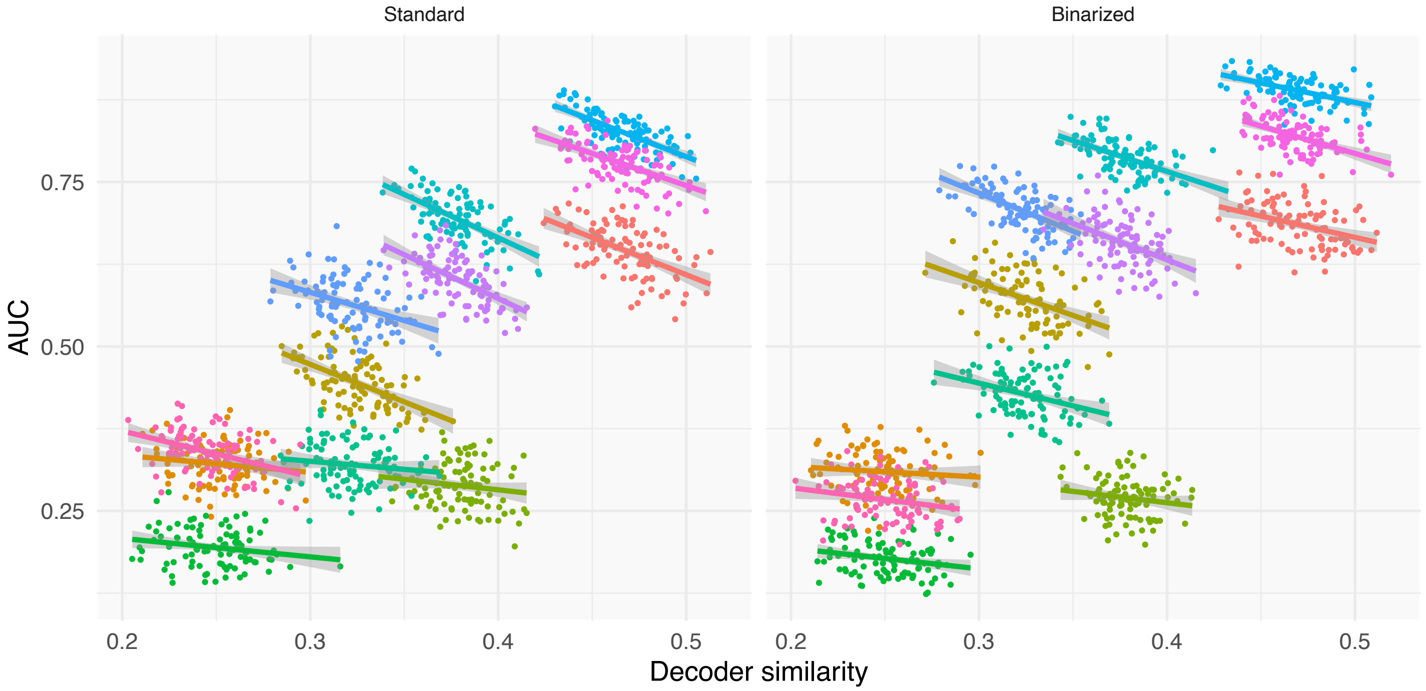


**Supplemental Figure 12. Location-only within-subject GLM decoders perform similarly to full GLM decoders.** To further assess whether location is the minimum necessary decoding function for successful decoding, we binarized the within-subject decoders from **Figure 4d** as a function of $z > 2.3$.

Left panel is the same as **Figure 6d-left panel**. It is reproduced here to enable direct comparison with location-only within-subject decoders performance.

Within-subject location-only decoders performed similarly to the full decoders regarding the relationship between decoder similarity and AUC. In this case, decoder similarity reflects the extent to which pairs of decoders were spatially coincident. Therefore, even in within-subject decoding, location information is the dominant parameter influencing performance.

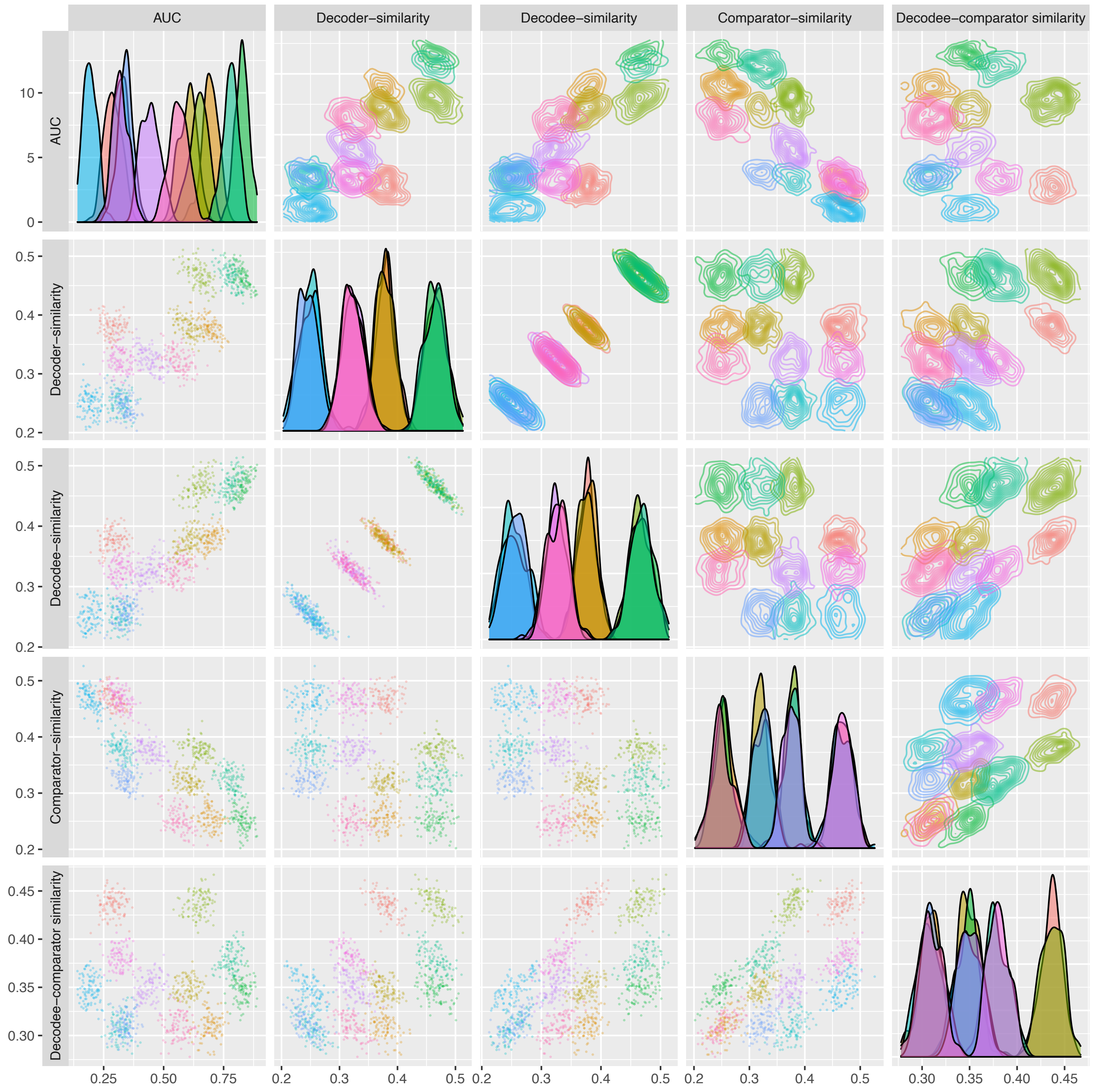


**Supplemental Figure 13. Determinants of within-subject discrimination performance for GLM decoders.** We examined pattern similarity within and between decoder, decodee, and comparator, to uncover determinants of performance. Data from Jimura et al. (*1*) were used to create and test within-subject decoders, for four cognitive tasks. Each decoder was created using the averaged GLM activity map from 6 of the 12 task repetitions, and tested on the remaining 6 (decodee) and 6 randomly selected replicates of a control task (comparators). Similarity of the *decoder* represents the average normalized dot product of the GLM maps used to create the decoder, and likewise for the similarity of the *decodee* and *comparator* (i.e., normalized dot products of the GLM maps in each sample). Decodee-comparator similarity is the average normalized dot product between all combinations of the decodee and comparator GLM maps. Note the positive correlations between AUC and decoder and decodee similarities (as shown in **fig. 4D left panel**), and the negative correlation between AUC and comparator similarity; however, there is no salient relationship between AUC and decodee-comparator similarity. Despite this, exploratory regressions found that the best, parsimonious multiple regressions for performance contained both decodee-similarity and decodee-comparator-similarity ($R^2 = 0.95$); the alternative multiple regression model contained comparator-similarity and decodee-comparator-similarity ($R^2 = 0.96$) (see **Supplementary Table 1**). Moreover,

multiple more complex models that are variants of these minimal models also account for most of the variance of decoding. Thus, task properties almost entirely determine within subject decoding performance.
Diagonal panels are density plots, off-diagonal panels are scatter-plots and corresponding contour plots.

**Supplementary Table 1. Similarity measures as a determinant of discrimination performance (AUC)**

| Model | Parameter | Within-subject Estimate ± SE | Within-subject Multiple $R^2$ | Between-subject Estimate ± SE | Between-subject Multiple $R^2$ |
|---|---|---|---|---|---|
| 1 | Intercept | 0.502 ± 0.004 | 0.61 | 0.598 ± 0.004 | 0.13 |
| | Decoder similarity | 1.96 ± 0.05 | | 0.84 ± 0.06 | |
| 2 | Intercept | 0.502 ± 0.003 | 0.66 | 0.598 ± 0.003 | 0.43 |
| | Decodee similarity | 2.09 ± 0.04 | | 1.50 ± 0.05 | |
| 3 | Intercept | 0.502 ± 0.004 | 0.58 | 0.598 ± 0.004 | 0.14 |
| | Comparator similarity | -1.93 ± 0.05 | | -0.87 ± 0.06 | |
| 4 | Intercept | 0.502 ± 0.006 | < 0.01 | 0.598 ± 0.005 | 0.01 |
| | Decodee-comparator similarity | 0.02 ± 0.13 | | 0.34 ± 0.09 | |
| 5 | Intercept | 0.502 ± 0.003 | 0.67 | 0.598 ± 0.003 | 0.51 |
| | Decoder similarity | 0.63 ± 0.10 | | 0.64 ± 0.05 | |
| | Decodee similarity | 1.5 ± 1.0 | | 1.4 ± 0.05 | |
| 6 | Intercept | 0.502 ± 0.002 | 0.90 | 0.598 ± 0.004 | 0.19 |
| | Decoder similarity | 1.51 ± 0.02 | | 0.57 ± 0.07 | |
| | Comparator similarity | -1.45 ± 0.02 | | -0.61 ± 0.07 | |
| 7 | Intercept | 0.502 ± 0.003 | 0.79 | 0.598 ± 0.004 | 0.17 |
| | Decoder similarity | 2.57 ± 0.04 | | 0.92 ± 0.06 | |
| | Decodee-comparator similarity | -2.23 ± 0.07 | | 0.58 ± 0.08 | |
| 8 | Intercept | 0.502 ± 0.001 | 0.94 | 0.598 ± 0.003 | 0.64 |
| | Decodee similarity | 1.63 ± 0.02 | | 1.63 ± 0.04 | |
| | Comparator similarity | -1.42 ± 0.02 | | -1.08 ± 0.04 | |
| 9 | Intercept | 0.502 ± 0.001 | 0.93 | 0.598 ± 0.003 | 0.67 |
| | Decodee similarity | 2.97 ± 0.02 | | 2.58 ± 0.05 | |
| | Decodee-comparator similarity | -2.82 ± 0.04 | | -2.14 ± 0.06 | |
| 10 | Intercept | 0.502 ± 0.002 | 0.83 | 0.598 ± 0.004 | 0.40 |
| | Comparator similarity | -2.78 ± 0.04 | | -2.04 ± 0.07 | |
| | Decodee-comparator similarity | 2.74 ± 0.06 | | 2.26 ± 0.10 | |
| 11 | Intercept | 0.502 ± 0.002 | 0.91 | 0.598 ± 0.004 | 0.42 |
| | Decoder similarity | 1.30 ± 0.04 | | 0.32 ± 0.06 | |
| | Comparator similarity | -1.69 ± 0.04 | | -1.84 ± 0.08 | |
| | Decodee-comparator similarity | 0.53 ± 0.09 | | 2.16 ± 0.10 | |
| 12 | Intercept | 0.502 ± 0.001 | 0.95 | 0.598 ± 0.003 | 0.68 |
| | Decodee similarity | 2.27 ± 0.04 | | 2.35 ± 0.07 | |
| | Comparator similarity | -0.77 ± 0.04 | | -0.34 ± 0.08 | |
| | Decodee-comparator similarity | -1.4 ± 0.09 | | -1.6 ± 0.1 | |
| 13 | Intercept | 0.502 ± 0.001 | 0.94 | 0.598 ± 0.003 | 0.68 |
| | Decoder similarity | 0.56 ± 0.04 | | 0.21 ± 0.04 | |
| | Decodee similarity | 2.4 ± 0.04 | | 2.5 ± 0.06 | |
| | Decodee-comparator similarity | -2.8 ± 0.04 | | -2.0 ± 0.08 | |
| 14 | Intercept | 0.502 ± 0.001 | 0.94 | 0.598 ± 0.003 | 0.68 |
| | Decoder similarity | 0.47 ± 0.04 | | 0.16 ± 0.05 | |
| | Decodee similarity | 1.2 ± 0.04 | | 1.6 ± 0.04 | |
| | Comparator similarity | -1.4 ± 0.02 | | -1.0 ± 0.05 | |
| 15 | Intercept | 0.502 ± 0.001 | 0.95 | 0.598 ± 0.003 | 0.68 |
| | Decoder similarity | 0.51 ± 0.04 | | 0.17 ± 0.04 | |
| | Decodee similarity | 1.84 ± 0.05 | | 2.31 ± 0.07 | |
| | Comparator similarity | -0.73 ± 0.04 | | -0.26 ± 0.08 | |
| | Decodee-comparator similarity | 1.5 ± 0.08 | | -1.6 ± 0.14 | |
| 16 | Intercept | 0.502 ± 0.002 | 0.91 | 0.598 ± 0.003 | 0.64 |
| | Decodee / Decodee-comparator | 1.02 ± 0.01 | | 1.16 ± 0.02 | |

We performed linear regression with AUC as the dependent variable and all possible linear combinations of similarity measures as the independent variables. All independent variables are mean-centered so that intercepts are interpretable and represent the estimate when all covariates are equal to the mean. Note, standard errors are influenced by the number of permutations and thus should not be used for inference.

In all models (where some of the variance is accounted for; all but model 4), our ability to capture discrimination performance is far superior when comparisons are done within subject, in contrast to between-subject comparisons.

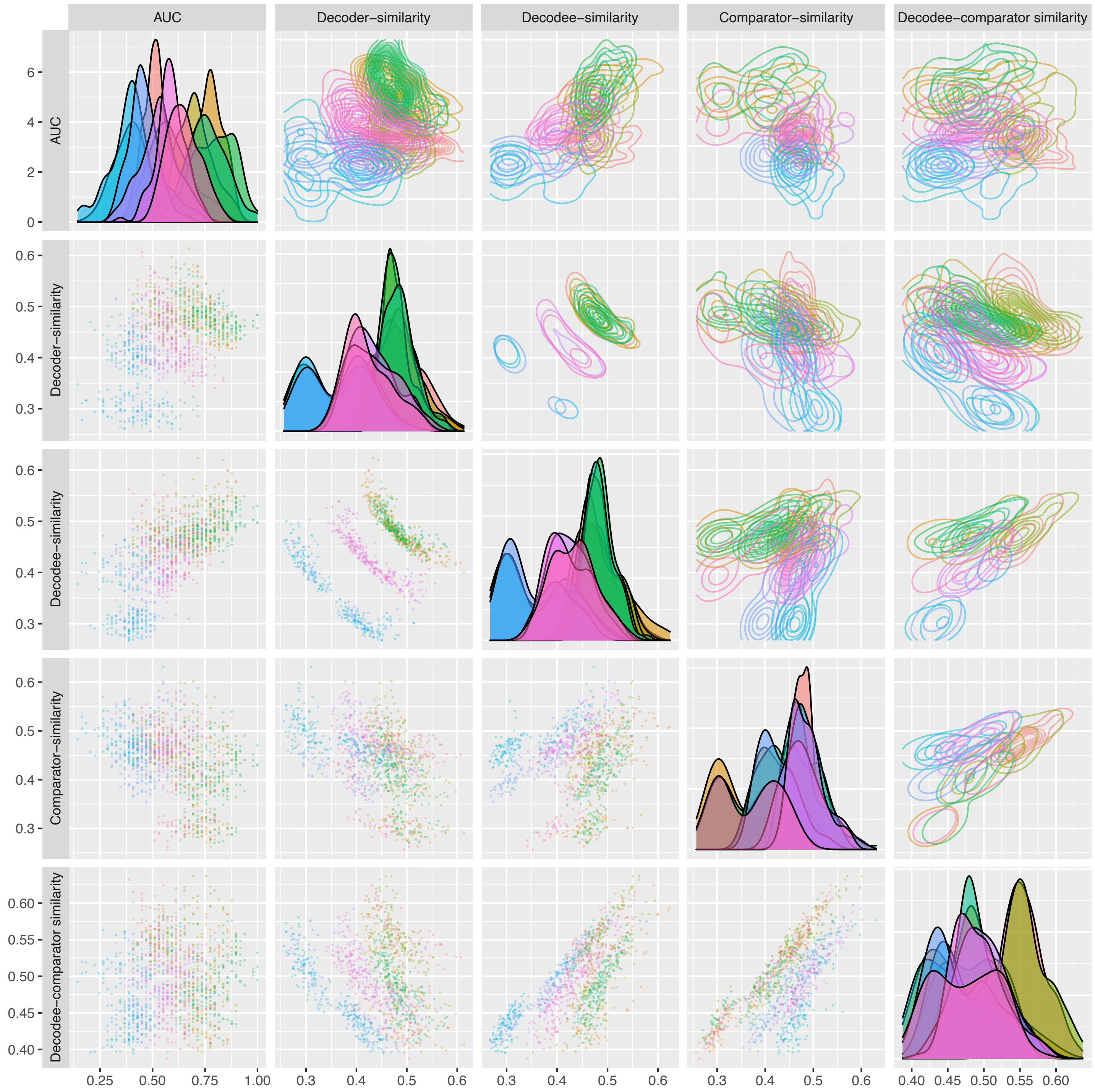


**Supplemental Figure 14. Determinants of between-subject discrimination performance for GLM decoders.** Data from Jimura et al. (*1*) were used to create and test between-subject decoders, for four cognitive tasks. Each decoder was created using the averaged GLM activity map from 6 of the 12 task repetitions, and it was tested on 6 other subjects for the corresponding task (decodee-s from different subjects) and 6 other subjects for a comparator task. Similarity of the *decoder* represents the average dot product of the GLM maps used to create the decoder, and likewise for the similarity of the *decodee* and *comparator* (i.e., dot products of the GLM maps in each sample). Decodee-comparator similarity is the average dot product between all combinations of the decodee and comparator GLM maps between different subjects. Note that all relationships are less clear than those in **Supplementary Figure S13**, which define relationships quantified in the regression analysis, shown in **Supplementary Table 1**.
